# Phototactic preference and its genetic basis in the planulae of the colonial Hydrozoan Hydractinia symbiolongicarpus

**DOI:** 10.1101/2024.03.28.585045

**Authors:** Sydney Birch, Lindy McGee, Curtis Provencher, Christine DeMio, David Plachetzki

## Abstract

**Background:** Marine organisms with sessile adults commonly possess motile larval stages that make settlement decisions based on integrating environmental sensory cues. Phototaxis, the movement toward or away from light, is a common behavioral characteristic of aquatic and marine metazoan larvae, and of algae, protists, and fungi. In cnidarians, behavioral genomic investigations of motile planulae larvae have been conducted in anthozoans (corals and sea anemones) and scyphozoans (true jellyfish), but such studies are presently lacking in hydrozoans. Here, we examined the behavioral genomics of phototaxis in planulae of the hydrozoan *Hydractinia symbiolongicarpus*.

**Results:** A behavioral phototaxis study of day 3 planulae indicated preferential phototaxis to green (523 nm) and blue (470 nm) wavelengths of light, but not red (625 nm) wavelengths. A developmental transcriptome study where planula larvae were collected from four developmental time points for RNA-seq revealed that many genes critical to the physiology and development of ciliary photosensory systems are dynamically expressed in planula development and correspond to the expression of phototactic behavior. Microscopical investigations using immunohistochemistry and *in situ* hybridization demonstrated that several transcripts with predicted function in photoreceptors, including cnidops class opsin, CNG ion channel, and CRX-like transcription factor, localize to ciliated bipolar sensory neurons of the aboral sensory neural plexus, which is associated with the direction of phototaxis and the site of settlement.

**Conclusions:** The phototactic preference displayed by planulae is consistent with the shallow sandy marine habitats they experience in nature. Our genomic investigations add further evidence of similarities between cnidops-mediated photoreceptors of hydrozoans and other cnidarians and ciliary photoreceptors as found in the eyes of humans and other bilaterians, suggesting aspects of their shared evolutionary history.

## Background

Light is a sensory cue of major importance in the marine environment, utilized by a vast diversity of organisms to tune biological processes. Light cues can dictate when to mate, movement through the water column, and in the case of sessile marine larvae, when and where to settle. Phototaxis, the movement toward or away from light, is a common behavioral characteristic of aquatic and marine metazoan larvae and of algae, protists, and fungi (Randel & Jékely, 2016).

*Hydractinia symbiolongicarpus* is a hydrozoan closely related to Hydra and other hydroids in the phylum Cnidaria, which occupies a key phylogenetic position in animal phylogeny as the evolutionary sister to Bilateria (2). The adult phase of *H. symbiolongicarpus* is sessile and resides on the shells of gastropods inhabited by hermit crabs in shallow sandy environments. The larval stage, known as a planula, is the only motile life history stage in this species. The spindle-shaped *H. symbiolongicarpus* planula becomes phototactic after day three of development (72 hours post fertilization), where larvae are competent to settle and undergo metamorphosis into a primary polyp (3). *H. symbiolongicarpus* planulae possess a nerve net that forms a neural plexus comprised of sensory neurons in the blunt, concave, aboral region of the larva (4,5). During settlement, planulae attach to the substrate at the aboral end, which is the hypothesized site of sensory integration. Much of the research into *H. symbiolongicarpus* planulae has focused on the induction of settlement by neuropeptides and chemical cues, where neuropeptides like RFamide play a role in relaying photosensory stimuli to epithelial muscular cells, directing the movement of the photo-response (6,7). However, the specificity of the phototactic response and its underlying sensory and developmental physiology are poorly understood.

Previous work on the sensory physiology of phototaxis in cnidarian planulae has largely focused on anthozoan species including anemones and corals. Coral planulae express multiple distinct opsin classes, indicating the possibility of multiple phototransduction systems (8,9). Additionally, planulae of different coral species may be phototactic to different wavelengths of light, including red wavelengths, suggesting a role for planula photosensitivity in ecological niche partitioning (10). However, anthozoan planulae possess several features that distinguish them from the planulae of hydrozoans and other medusozoans. First, comparative analyses indicate that anthozoan genomes house an expanded diversity of opsin genes that includes the cnidarian-specific opsin classes cnidops and anthozoan-specific opsin (ASO), in addition to ciliary opsin and xenopsins that are also present in bilaterians (8,9,11–14). In contrast, the opsin palate of hydrozoan genomes examined thus far is restricted to cnidops class genes (12,14–16). In addition, anthozoan planulae of the order Actinaria, which includes anemones, differ morphologically from other cnidarian planulae in their possession of a distinct apical tuft organ located on the aboral end that is thought to be sensory in nature (17–21).

Here we investigate the photosensory preference of the *H. symbiolongicarpus* planula and explore the genetic basis of this behavioral response using phototaxis assays, systems-level comparative developmental transcriptome analyses, and microscopical investigations of gene expression. We report that *H. symbiolongicarpus* planulae are strongly phototactic to green and blue wavelengths of light, but not red wavelengths. RNA-seq analyses reveal a significant increase in gene expression of photosensory genes in larval stages that precede phototaxis, including those involved in phototransduction, such as opsin, structural components of photoreceptors, and several developmental factors that have not previously been reported in cnidarian species. IHC and RNA FISH studies revealed that opsin transcripts localize to bipolar photoreceptor neurons that occupy the aboral neural plexus of the planula. Surprisingly, we identify a cnidarian homolog of transcription factor CRX, which is central to rod/cone fate determination in mammals, that is also strongly expressed in *H. symbiolongicarpus* photoreceptors. Together, this study reveals that *H. symbiolongicarpus* planulae are differentially capable of perceiving various wavelengths of light, and the photoreceptors upon which this behavior is based may utilize an ancient molecular toolkit related to the ciliary photoreceptors of bilaterian animals. Our results diverge from recent findings in anthozoan planulae, and the molecular and cellular composition of apical tuft organs (19).

## Methods

### Field Collection of *H. symbiolongicarpus*

Colonies of *H. symbiolongicarpus* affixed to hermit crabs (*Pagurus longicarpus*) were collected at Barnstable Bay, MA (41°42’39’’N 70°16/34’’W) in the spring of 2021. The colonies were cultured and maintained according to Plickert & Schneider (2004) and Frank, Nicotra, & Schnitzler (2020) where they were kept in an incubator on a 12/12 L:D cycle at 18°C. Spawning was induced by placing a single male and a single female colony under bright light in a dissecting bowl filled with filtered seawater at room temperature. This was done in three replicates using haphazardly chosen male and female colonies. For each spawning, planulae were collected and placed in petri dishes in filtered seawater. Planulae were collected for sequencing at 24-hour intervals over four days. Phototaxis experiments and microscopy were conducted on day 3 planulae.

### Larval phototaxis study

Phototaxis was studied in planulae larvae on the third day of development (72 hours post fertilization) when larvae are known to respond to light cues (22). We used a behavioral apparatus consisting of a raised piece of black acrylic with three 1/8-inch holes spaced 8 inches apart. Below the acrylic, we placed three small, heatless LED lights tuned to the wavelengths of 470nm, 523nm, and 625nm (SuperBright LEDs: RL5-B12120; RL5-G16120; RL5-R12120 respectively) into the holes. A petri dish containing 25mL of filtered seawater was placed above each LED light (Fig. 1A). Experiments were conducted in total darkness. Each replicate examined the three wavelengths of light using three Petri dishes. We performed five replicates. For each experiment, 100 larvae were added to each petri dish where larvae were spatially randomized with a pipette before the start of the experiment when ‘before’ photographs were taken (Fig. 1A). The three light stimuli were then applied for eight hours, and an ‘after’ photograph was taken. Phototaxis was assessed by comparing the number of larvae that had migrated within 1 cm of the LED light to the number of larvae that had not. Experiments were conducted across three independent genetic crosses from wild-caught individuals. Welch two-sample t-tests were implemented for each wavelength to statistically interrogate the number of larvae that responded to the light stimulus.

### Library preparation, sequencing, and read processing

We collected three replicates of 15-20 larvae for each day of development (days 1-4) and extracted total RNA using the PureLink RNA Mini Kit (Thermo Fisher Scientific; Cat no. 12183018A) according to the manufacturer’s instructions. Libraries were created using 1000ng of total RNA with the NEBNext Ultra 2 Directional RNA Library Kit following the Poly(A) mRNA Magnetic Isolation Module (NEB #E7490) and sequenced on an Illumina Hi-Seq 2000. Reads were processed following a previously established pipeline (23) and assembled using the Oyster River Protocol (ORP), which performs read processing (i.e. read trimming, read normalization, read error correction) and uses multi-kmer and multi-assembler approaches to generate the reference assembly with TransRate and BUSCO quality metrics (24).

### Gene expression analysis

We used Salmon to quantify transcripts (Patro et al., 2017) and EdgeR to estimate differential gene expression between all pair-wise comparisons of the four developmental days (26). Differentially expressed genes were identified at a p-value of 0.0001 using the Benjamin-Hochberg correction. TransDecoder was used to translate transcripts into proteins (27), and cd-hit (28) was used to reduce the number of duplicate protein models. The reduced *H. symbiolongicarpus* protein models were used in OrthoFinder (29) with the following taxa from publicly available data: *Homo sapiens* (30), *Drosophila melanogaster* (31), *Hydra vulgaris* (https://research.nhgri.nih.gov/hydra/), *H. symbiolongicarpus* (https://research.nhgri.nih.gov/hydractinia/), *Ectopleura crocea* (23), and *Nematostella vectensis* (32). Orthogroups were annotated based on human sequences that were present in gene sets obtained from the Gene Set Enrichment Analysis (GSEA) https://www.gsea-msigdb.org/gsea/index.jsp (33–35), including GO_Sensory_Perception_Of_Light_Stimulus and GO_Sensory_System_Development. We annotated *H. symbiolongicarpus* transcripts based on their shared occupancy in orthogroups with human genes from the above gene sets. Differentially expressed genes identified from each gene set were analyzed using the STRING database which includes known and predicted protein-protein interactions (36). For each gene set, we examined significantly upregulated orthogroup gene symbols across developmental days, per day, and combined, and created interaction networks using selected Gene Ontology (GO) terms or Annotated Keywords (UniProt) of interest as annotations. In addition to the significantly differentially expressed genes, we analyzed all genes expressed in the gene sets that were not significantly differentially expressed, and the orthogroups present in gene sets that did not contain *H. symbiolongicarpus* transcripts. All scripts and workflows can be found on GitHub: https://github.com/sydney-birch/Hydractinia_phototaxis.

### Immunohistochemistry, RNA fluorescent *in situ* hybridization (FISH), and confocal microscopy

Planulae were fixed in seawater with 4% paraformaldehyde overnight at 4°C. For immunohistochemistry, planulae were washed five times with five-minute incubations in Phosphate Buffered Saline plus 0.01% Tween 20 (PBST), then blocked in PBST plus 20% normal goat serum (NGS) for two hours. Primary antibodies: mouse monoclonal anti-α-Tubulin (Sigma T8203), and rabbit polyclonal anti-FMRF amide (EDM Millipore, AB15348), were added overnight at 4°C (1:500 in PBST plus 20% NGS). Primary antibodies were removed, and planulae were washed five times with five-minute incubations in PBST. Secondary antibodies: AlexaFlour™ 546 goat anti-mouse (Invitrogen, A11030), AlexaFlour™ 633 goat anti-rabbit (Invitrogen, A21071), and Alexa Fluor™ 488 Phalloidin (1:20), were added overnight at 4°C (1:500 in PBST plus 20% NGS). Planulae were washed five times with five-minute incubations in PBST and mounted in Prolong™ gold antifade plus DAPI (Invitrogen, P36935).

For RNA fluorescent *in situ* hybridization, we followed the protocol from (23) for designing RNA FISH probes of the highest expressed transcripts of interest and for staining. RNA FISH probes were designed using the Stellaris RNA FISH platform (Biosearch Technologies) with the custom probe design service. Cnidopsin (Hs_t.88569) riboprobe sets were labeled with FAM (488 nm), CNG ion channel (Hs_t.35573) riboprobe sets were labeled with TAMARA (561 nm), and CRX (Hs_t.47300) riboprobe sets were labeled with Quasar670 (635 nm). For samples treated with FISH probes and IHC antibodies (Fig. 7), we first performed the *in situ* hybridization protocol detailed in (23), then refixed samples in 4% PF in PBST for 30 minutes at room temperature. Samples were washed five times with five-minute incubations in PBST and then incubated overnight in primary antibody solution containing anti-acetylated alpha-tubulin (1:500; Sigma, T6793) in PBST plus 20% NGS. Samples were washed five times with five-minute incubations in PBST and then incubated with secondary antibody (Alexa Flour 546-conjugated anti-mouse IgG, 1:1000; Life Technologies, A11030) overnight at 4°C. Samples were washed five times with five-minute incubations in PBST and then mounted in ProLong antifade mountant with DAPI (ThermoFisher, P36941). Samples were imaged on a Nikon A1R HD confocal microscope.

## Results

### Planulae are strongly phototactic by developmental day three to green and blue wavelengths of light

We examined the phototactic behavioral response of motile, day 3 (72 hours post-fertilization) *H. symbiolongicarpus* planulae to blue (470nm), green (523 nm), and red (625 nm) wavelengths of light by assessing the number of larvae that migrated to within 1 cm of an LED affixed to the underside of a transparent petri dish after eight hours. Planulae were strongly phototactic towards green (*p*=0.01), and blue light (*p*=0.0015). Conversely, larvae were not phototactically drawn to red wavelengths of light (*p*=0.66). Images taken after such experiments show planulae tightly clustered around green and blue LEDs (Fig 1). In contrast, in the presence of red light, planulae remained randomly distributed with respect to the light stimulus after eight hours.

### Developmental transcriptomics and orthology analyses

We investigated the genetic basis for phototaxis in *H. symbiolongicarpus* planula larvae by performing a developmental transcriptome study. Here RNA-seq data were derived from three replicates of 15-20 pooled larvae that resulted from a single mating of wild-caught colonies, collected at each developmental day (1–4). In *H. symbiolongicarpus*, day-three larvae display phototaxis, and day-four larvae are competent to settle and metamorphose into a primary polyp (5,6,22,37). Illumina RNA-seq data included over 298 million reads across three replicates of each developmental day, with an average of 24 million reads per replicate. A reference transcriptome was assembled and contained 92,784 sequences, with a TransRate score of 0.39, and a BUSCO score of 100. Multidimensional scaling (MDS) of quantitative estimates of transcript abundance clustered RNA-seq datasets by developmental day (Fig 2A). Pairwise comparisons indicated the greatest total expression differences between developmental days one and four, and the weakest total expression differences between developmental days two and three (Fig 2B).

The genetic basis for phototaxis in *H. symbiolongicarpus* planulae was examined by interrogating orthogroups produced from orthology analyses (29) using previously derived functional gene sets from the Gene Set Enrichment Analysis (GSEA) database (33–35). Our approach links specific *H. symbiolongicarpus* transcripts to human gene symbols based on their shared occupancy in formally defined orthogroups, which represent groups of homologous genes derived from a shared last common ancestor (23,29,33,38). Orthology analyses utilized a dataset including the present data plus an additional five genome-scale protein coding datasets from cnidarian and bilaterian species, including human.

Opsin-mediated photobehavior has been described in cnidarians previously (8,9,44,12,16,23,39–43). We, therefore, identified between-day differentially expressed transcripts with membership in GSEA gene sets that contained opsin, which included the *sensory perception of light* (Fig 3) and *sensory systems development* (Fig 4) gene sets, the latter including genes involved in mechanoreception and chemoreception as well.

### Gene expression during planula development mirrors aspects of ciliary photoreceptor development and function

Analyses over development of significantly differentially expressed genes present in the *sensory perception of light stimulus* gene set revealed a pattern of congruence between various aspects of vertebrate retina development and function, and *H. symbiolongicarpus* planula development and the expression of phototaxis (Fig 3). We depict our findings using STRING analyses, which display graphical linkages between differentially expressed transcripts based on databases of known protein-protein interactions (45). From the perspective of the *sensory perception of light stimulus* gene set, Day 1 of planula development is characterized by the expression of regulatory factors like SOX and POU transcription factors, nuclear receptors NR2E1 and NR2E3, and leucine zipper transcription factor NRL (46). Also at peak expression in Day 1 planulae are signaling components including transducins GNAT1 and GNAT2 (Fig 3C). A shift in the transcriptomic repertoire, from neurogenesis and development on Day 1, to photoreceptor maintenance and sensory physiology, occurs on Day 2 (Fig 3C) where we observe peak transcript expression for opsin, multiple semaphorin transcripts (SEMA5B), photoreceptor-specific EF-hand Calcium-binding protein (PPEF2), and multiple Major Intrinsic Protein (MIP) transcripts. Day 3 differentially expressed planula transcripts include a single *H. symbiolongicarpus* transcript homolog of CRX, VSX, and RAX transcription factor paralogs known from mammalian photoreceptor development (47). Genes involved in photoreceptor structure and function also have peak expression on Day 3 including adhesion GPCR V1 (ADGRV1) and guanylyl cyclase (GUCY2F, GUCY2D). Day 4 differentially upregulated planula transcripts are mostly associated with structural components including the stereociliary scaffolding proteins usherin (USH1C) and CDH23, of which *H. symbiolongicarpus* has multiple transcripts (Fig 3), and myosin III (MYO3B, MYO3A). In addition, we note strong expression of delta (DLL4) in Day 4, which, together with notch, which is not present in our transcriptome dataset, has been implicated in repressing cone photoreceptor state (48). Our analysis reveals a transition from early genes involved in neurogenesis and proliferation (Figure 3C), to more specialized regulatory, structural, and physiological features of ciliary photoreceptors in later developmental days (Fig 3C). Protein-protein interaction (PPI) p-values are significant for each developmental day, which is expected given that our analyses are constrained by gene sets with known function. However, PPI enrichment is most significant in developmental days 3 and 4.

Transcripts that are differentially expressed between days may be the basis for the developmental dynamics observed in planula development. However, other genes, known to be involved in the *sensory perception of light* gene set are also expressed in our transcriptome data but are not significantly differentially expressed between days (supplemental figure 1), perhaps due to the nature of our bulk RNA-seq dataset. Transcripts of this type include homologs of alpha, gamma, and zeta lens crystallin (CRYGA, CRYGC, CRYGB, and CRYZ), cyclic nucleotide-gated ion channel (CNGA3, CNGB3), and phosphodiesterase (PDE). We also compared the functional interactions of significantly differentially expressed transcripts to expressed transcripts using enrichment analyses (supplemental figure 1) and found that all the functional categories involved in development and phototransduction are enriched in the significantly differentially expressed gene set.

It is also informative to examine components of the *sensory perception of light* gene set that are not present in our dataset (supplemental figure 2). Here, we note several accessory components that act to modulate phototransduction that are not present in our *H. symbiolongicarpus* developmental transcriptome dataset, including RGS9 and RGS16, GRK7, RGR, and PDE6G among others. Expectedly, several structural genes associated with the lens are also missing in our transcriptome data including the lens-associated proteoglycan keratocan (KERA), the beta crystallins (CRYBB1, CRYBB2, CRYBB3, CRYBB4), and the lens-specific gap junction proteins (GJA8 and GJA3), aligning with the lack of a lens structure in this species.

### Limited correspondence between planula development and other bilaterian sensory systems

Having identified much overlap between a gene set based on the development, structure, and physiology of ciliary photoreceptors, we next sought to understand what homological linkages might exist between planula development and the other senses. We examined the *sensory systems development* gene set, which includes genes involved in the regulation of the development of vision and photosensitivity, taste and olfaction, and hearing and mechanosensitivity (33) (Fig 4). However, only genes with pleiotropic function in photosensitivity and other sensory modalities (GNB1, GNAT1, GNAT2, USH1C, TBX2) were recovered from this analysis. In contrast to the *sensory system development* gene set, PPI enrichment p-values are most significant at developmental day 1 for the *sensory systems development* gene set.

As with the previous gene set, many of the transcripts expressed during planula development have homologs in the *sensory system development* gene set but are not differentially expressed. These include homologs of WNT, ATOH7, MITF, and others (supplemental Fig. 3). For comparison, genes not expressed by *H. symbiolongicarpus* larvae in the *sensory system development* gene set is given in supplemental Fig 4., and genes from the total gene set of transcription factors is given in supplemental Fig. 5.

### An aboral sensory array is present in phototactic planulae

The cellular and morphological basis for larval phototaxis in *H. symbiolongicarpus* planulae was investigated using immunohistochemistry and confocal microscopy (Fig 5). By developmental day 3, planulae possess an aboral plexus of anti-acetylated-tubulin-reactive sensory neurons that correspond to the site of sensory integration and settlement, as previously described (5,7,49,50) (red; Fig 5, inset, and supplemental video 1). However, FMRFamide immunoreactivity (magenta; Fig 5) is most intense in the oral end of the planula, the site of the future mouth in the primary polyp. Ordered musculature, as revealed by phalloidin staining (green; Fig 5), is mostly confined to the middle latitudinal portion of planulae. Prominent stereo ciliary projections associated with cnidocytes are mostly confined to the oral end but are also associated with cnidops-expressing photoreceptors in the aboral end of the planula, where fewer cnidocytes were observed (Figs 5 and 6). The oral end of the planula resembles an adult tentacle in morphology (Fig 5 and supplemental video 2).

### Immunohistochemistry and in situ hybridization reveal an opsin-expressing aboral sensory neural plexus

We examined the structure of *H. symbiolongicarpus* planulae using immunohistochemistry, fluorescent in situ hybridization, and confocal microscopy. We first examined the expression of opsin transcript Hs_t.88569 together with immunohistochemical staining with anti-acetylated Tubulin (Fig 6). Hs_t.88569 is the most prominently expressed opsin transcript in our dataset (Figs 3 and 4) and is a member of the cnidops family of opsins (15) (supplemental figure 6). As expected, Hs_t.88569 transcripts strongly localize to the cilia of bipolar sensory neurons that comprise the aboral neural plexus (Fig 6B). *H. symbiolongicarpus* transcripts localize specifically to ciliary projections of apical neurons that we interpret as photoreceptors. Optical sections also reveal a subnetwork of neurites connecting bipolar sensory neurons, which form an array around the aboral end of the planula.

### Ciliary-like gene expression in planula photoreceptors

In addition to cnidopsin (Hs_t.88569), we also examined the expression of other transcripts including CNG ion channel (Hs_t.35573) and CRX (Hs_t.47300) that were detected in our transcriptome screens, using multiplex fluorescent in situ hybridization (Fig 7). Homologs of each of these transcripts have been implicated in vertebrate photoreceptor development and function (51,52), and CNG ion channels are thought to function in cnidops-mediated phototransduction (16,39,40). All transcripts showed elevated staining in the aboral region, the site of settlement, with little detectable staining in the oral region. Higher resolution examination reveals that each of these transcripts localize specifically to opsin-expressing bipolar photoreceptor neurons that comprise the aboral sensory plexus (Fig. 7 A and B).

## Discussion

### Phototaxis and dispersal in cnidarian planulae

Larval phototaxis is widespread in the marine environment and has been implicated as a strategy for predator avoidance, dispersal, diel vertical migrations, and larval settlement (1,53–56). *H. symbiolongicarpus* planulae are phototactic and neuropeptides including RFamide play a role in relaying photosensory stimuli to epithelial muscular cells, directing the movement of the photoresponse (6,7). Our phototaxis experiment demonstrates that *H. symbiolongicarpus* planulae respond specifically to green and blue wavelengths of light, where larvae have the strongest sensitivity to green wavelengths (Fig 1). This phototactic response differs from anthozoan planulae that have been examined. Depending on the species, coral planulae may be positively phototactic to red wavelengths of light (41), or other wavelengths (10), indicating that they are tuned to light cues that provide information about species-specific niches (57). Together with previous studies, hydrozoans seem to show the strongest sensitivity to light in the blue-green spectrum irrespective of life history stage (16,23,58).

Phototaxis in *H. symbiolongicarpus* has been linked to identifying a substrate for attachment. *H. symbiolongicarpus* colonies typically reside on gastropod shells inhabited by hermit crabs which are found in shallow, sandy environments (37). Larvae attach to hermit crab shells by first attaching to an elevated structure (grain of sand or a stone) with their aboral blunt end. They then are thought to wait for an object to pass them and attach at their tapered oral end using atrichous isorhizas and desmoneme nematocytes (37). Vertical-facing surfaces are predominately illuminated by blue-green light (10). Therefore, the photo behavior observed here is consistent with larvae seeking suitable surfaces for which to capture moving hermit crab targets for settlement. Additionally, green sensitivity is also consistent with larvae migrating to the shallow depths where hermit crabs reside.

In *Hydractinia echinata*, a species closely related to *H. symbiolongicarpus* where the adult stage also resides on hermit crab shells and planulae are also phototactic, phototaxis was interrupted in the presence of hermit crabs (59). Thus, a sensory mechanism other than phototaxis, likely involving the perception of chemical and/or mechanical cues, is required for settlement on the proper substrate. Similarly, we recently described the interaction between light, chemical, and mechanical cues in larval settlement in another hydrozoan species *Ectopleura crocea* (23), furthering the idea that the integration of multiple sensory cues may be a common mechanism for larval settlement (60–62).

### Developmental transcriptomics implicate photosensation and set the stage for metamorphosis

Our research strategy uses previously determined gene sets and developmental transcriptome analyses as a lens to explore sensory function in developing planulae. This approach is likely to identify copious overlap between differentially expressed genes and genes from selected sensory gene sets because many of the underlying processes including neurogenesis and cell proliferation, are present in planula development; and cnidarian genomes share many cellular and genetic homologs with vertebrates and other model species (63,64), from which functional gene sets were obtained (33). While the simple feature of being differentially expressed between days does not directly implicate such transcripts in phototaxis or sensory function, the observed patterns suggest hypotheses on the genetic modules associated with phototaxis in *H. symbiolongicarpus*, and their possible evolutionary histories.

Several homologs of key regulators of retinal progenitor cells and retinal cell type specification are differentially expressed in planula development (Fig. 3). For example, proliferation in retinal progenitor cells and early neurogenesis in planulae both utilize SOX transcription factors. In vertebrates, SOX transcription factors are also important regulators of retinal progenitor cell differentiation in developing retinae, where different paralogs influence proliferation and cell fate determination in a diversity of retinal cell types, including rods and cone photoreceptors (65). Cnidarian SOXB transcription factors have been previously investigated in *H. symbiolongicarpus* (66) and in the anthozoan *Nematostella vectensis* (67) where SOXB helps specify neural progenitors and their differentiation, however planulae have not been investigated. POU transcription factors are also identified in our analyses. POU transcription factors cooperate to control cone photoreceptor specification in developing mouse retinae, and inactivation of human paralog Pou2f2 which leads to the expression of rod-inducing NRL (68). Our analyses indicate that both POU and NRL peak in expression in day 1 planulae. POU transcription factors are ancient (69) and have been implicated in stem cell regulation in *H. symbiolongicarpus* (70) and in the specification of hair cells in *N. vectensis* (71). Also highly differentially expressed in day 1 planulae are single transcripts corresponding to human paralogs NR2E1 and NR2E3, and CRX, VSX, and RAX (Fig 3), respectively. In mammalian retinae, the nuclear receptor NR2E3 works with the cone-rod homeobox protein CRX and NRL to activate rod-specific gene expression (72,73). Therefore, the core components of the rod-cone differentiation network (47) are among the differentially expressed genes in planulae development in *H. symbiolongicarpus*.

Genes involved in photoreceptor structure and function peak in their expression at developmental day 2, prior to the onset of phototaxis. Such genes include cadherin (CDH23) and usherin (USH1C), which function as tip links (74) and ankle links, respectively, in stereocilia. Both proteins are structural components of photoreceptors (74–76) and mechanoreceptive hair cells in mammals (77). In cnidarians, CDH23 (74,78,79) and USCH1C (80) have been investigated in *N. vectensis*, where they are expressed in hair cells. In addition, we recovered multiple semaphorin transcripts involved in axon guidance (SEMA5B) (81), transducins GNAT1 and GNAT2 (82), and opsin of the cnidops family (14,15). Adhesion GPCR V1 (ADGRV1), associated with nervous system development (83), and myosin III (MYO3B, MYO3A) an actin-based motor protein with kinase activity present in photoreceptors and stereocilia in general, are also differentially expressed (84–86). The involvement of structural and physiological loci in the development of phototactic planulae suggests that many of these components were present in ciliated photosensory, or multisensory neurons in the ancestor of cnidarians and bilaterians (23,87). Our STRING analyses also show that protein-protein interactions are most enriched in developmental days 3 and 4, consistent with the expression of phototaxis behavior, and that the foundation for sensory integration is laid early in development.

Some genes that have been suggested to play a role in phototransduction, but presently lack conclusive functional evidence are also among the differentially expressed genes, including PPEF2, the serine/threonine phosphatase/EF-hand calcium-binding protein (88). Interestingly, this gene is homologous to the retinal degeneration C (rdgC) gene from *Drosophila melanogaster* where it may function in the Ca+-dependent modulation of phototransduction by catalyzing the dephosphorylation of opsin (89,90). The presence of homologs of PPEF2 in ciliary, rhabdomeric, and possibly cnidops photoreceptors would indicate an ancestral mechanism for modulating photoreception that may have predated the split between cnidarians and bilaterians and also the divergence of rhabdomeric and ciliary phototransduction cascades and cell types (13).

It is also interesting to consider the genes present in *the sensory perception of light* gene set that were not recovered in our analyses. Here, missing are a set of genes that function to attenuate phototransduction including RGS9 and RGS16 (91,92), GTPase activating proteins that accelerate the deactivation of G proteins (91), GRK7, a GPCR kinase involved in the deactivation of opsin signaling (93), and RGR, the retinal G protein receptor and putative photoisomerase (94). The existence of a cnidarian photoisomerase was interpreted based on immunohistochemistry data in the camera eyes of the box jelly *Tripedalia* (95), but RGR retinochromes of the type present in deuterostomes and protostomes are so far absent in cnidarian genomes (11).

The current model for cnidops-mediated phototransduction begins with the activation of a G_s_ G-protein, which activates adenylate cyclase (AC), which in turn catalyzes the production of cAMP, leading to the opening a CNG ion channel (9,16,39,40,96,97). The gene sets used in the annotation of orthogroups were derived from studies in mostly vertebrate model organisms, which use a different mode of ciliary phototransduction. Therefore not all components of cnidops-mediated phototransduction are present in the gene set annotations we used to interrogate our data. However, additional searches revealed AC expression in our transcriptome data. Ciliary phototransduction in mammals is driven by the activation of the G-protein transducin (Gt), which activates phosphodiesterase (PDE6). Activated PDE6 hydrolyzes the secondary messenger cGMP to non-cyclic conformations, which leads to the closing of CNG ion channels (51,98). The activation of PDE6 by Gt G-proteins involves the displacement of its subunit, PDE6 gamma (PDE6G), an intrinsically disordered protein that remains bound to PDE6 while inactive. The rapid release of inhibition of PDE6 by PDE6G is thought to provide the fast cascade dynamics required for image-forming vision in the mammalian retina (99), but this is an unlikely requirement of most cnidarian photosystems including *H. symbiolongicarpus* planulae (53,100). In fact, cnidops-mediated phototransduction bears similarity to olfactory transduction in mammals, where cascade dynamics are much slower than ciliary photoreceptor dynamics (39). PDE6 gamma was not recovered in our data but was weakly suggested in earlier analyses of the eyes of the box jelly *Tripedalia crystophora* (101).

### Other sensory systems

In addition to phototaxis, *H. symbiolongicarpus* planulae also integrate information from the chemical and mechanical environments (102) into settlement behavior. We screened differentially expressed planula transcripts using the *sensory systems development* gene set (33), which includes regulatory factors involved in the development of vision and photosensitivity, taste and olfaction, and hearing and mechanosensitivity (Fig 4). Annotations of differentially expressed transcripts throughout the developmental time course mostly represent neurogenesis, sensory system development, and specific photoreceptor functions, and show much overlap with the *sensory perception of light* gene set (Fig 3). Most of the genes recovered by this analysis are pleiotropic across sensory systems including the G-protein beta subunits GNB1, GNAT1, and GNAT2, which are involved in the perception of both light and chemical cues. Similarly, USH1C is a component of both mechanoreceptors and photoreceptors (86,103). In mammals, TBX2 is best known as a master regulator of inner vs. outer hair cell specification (104), however, it was recently identified as a factor controlling cone photoreceptor cell differentiation in *Danio rerio* as well (105). The lack of overlap between the differentially expressed genes in planula and non-photosensory modalities in the *sensory systems development* gene set may reflect both, a lack of conservation in modes of chemo-and mechano-sensation between cnidarians and mammals, and our lack of understanding of the functional components of these modalities. In contrast to the *sensory perception of light* gene set, PPI enrichment p-values are highest in day 1 for the *sensory systems development* gene set, suggesting that most of its correspondence with *H. symbiolongicarpus* planula development involves protein-protein interactions involved in foundational developmental processes.

In addition, many genes included in the *sensory systems development* gene set are of general importance in nervous system development and are expressed during *H. symbiolongicarpus* planula development, but not significantly differentially expressed across days. Often, such genes are pleiotropic and may be associated with non-sensory-cell-specific expression. Interestingly, functional categories that are significantly enriched among differentially expressed transcripts, compared to transcripts that are expressed but not differentially so, include phototransduction, sensory perception of light stimulus, and sensory perception of taste, the latter of which is driven by the expression of transducins, which function in both photoreception and olfaction (Fig 4; supplemental figure 3).

### The aboral sensory array as the site of sensory integration

Our bulk transcriptome analyses highlight the shared expression of transcripts involved in planula development in *H. symbiolongicarpus* planulae and ciliary photoreceptor development and function. We used fluorescent *in situ* hybridization to examine whether this functional overlap is constrained spatially in planulae, or in a cell type-specific manner. By developmental day 3, planulae show an aboral density of neural cells that are stained by the anti-acetylated tubulin antibody (Figs 5-6). In contrast, most FMRFamide immunoreactivity is confined to the oral end of planulae, and to a subpopulation of neurons (Fig 5). In *H. symbiolongicarpus*, the oral end of the planula is associated with the future mouth of the primary polyp. Thus, the FMRFamide immunoreactivity we observe could be associated with neurons destined for the adult nervous system. FMRFamide is also associated with a subpopulation of sensory neurons in the planulae of *N. vectensis* (106) and the scyphozoan *Aurelia aurita* (107), and in both cases is confined to the aboral end of the planulae, in contrast to our observations in *H. symbiolongicarpus*.

The aboral end of *H. symbiolongicarpus* has long been viewed as the site of sensory integration (37), however, our data add new insights into its structure and function. First, our microscopical data indicate greater neuronal organization in this region than previously described, indicating an array of bipolar sensory neurons that project outward across the entire aboral region, clearly visible in cross-sections (Figs 5-7, supplemental video 1). This bipolar neuron array is also connected by ganglion cells (Fig 6). Suggestions of this aboral neural plexus were indicated previously (4,5,114,6,49,108–113). Second, aboral bipolar sensory neurons express cnidops-class opsin and other genes involved in ciliary phototransduction and photoreceptors (e.g., CNG ion channels, opsin, and CRX-like transcription factors) (Fig 6 and 7). The expression domains of these ciliary photoreceptor transcripts roughly localize to the aboral half of the planulae and are strongly co-expressed in the ciliary regions of bipolar neurons (Fig 6 and 7). Subcellular localization of functional transcripts including opsins has also been described in larval photoreceptors of another hydrozoan species, *Ectopleura crocea* (23), and in mammalian cones (115). Together, our data indicate that the aboral end of planulae corresponds to a developmental field enriched for photoreceptor structure and function, but the expression of other sensory modalities in this region remains to be explored.

## Conclusions

Light is a sensory cue of major importance in the marine environment, utilized by a vast diversity of organisms to tune biological processes. We combined a larval phototaxis study with developmental transcriptomics, immunohistochemistry, and *in situ* hybridization to better understand the molecular determinants of phototaxis in *H. symbiolongicarpus* planulae. Analyses of differentially expressed transcripts indicated much overlap between functional modules associated with ciliary photoreceptors including development, morphogenesis, and phototransduction, and the onset of phototaxis in *H. symbiolongicarpus* planulae. Shared utilization of components of the rod/cone developmental differentiation network in both hydrozoan planulae and vertebrate retinae suggests it may have evolved prior to the split between cnidarians and bilaterians. Portions of gene sets that are not captured in our dataset also provide clues on the evolution of more specialized and sensitive photosensory systems in bilaterians. In addition, microscopical investigations reveal the aboral sensory neural plexus as the site of photosensory transcript expression making it the likely site of sensory integration. Our approach is sufficient for generating hypotheses on the determinants of planula development and sensory function and offers a useful comparison to the motile larvae of other species.

## List of Abbreviations

hpf: Hours post fertilization
DEGs: differentially expressed genes
IHC: Immunohistochemistry
FISH: fluorescent in situ hybridization

## Declarations

### Availability of data and materials

The datasets supporting the conclusions of this article are available in the SRA database, Accession: PRJNA1046120; and in Dryad, https://doi.org/10.5061/dryad.s1rn8pkfg. Source code can be found in the following GitHub, https://github.com/sydney-birch/Hydractinia_phototaxis.

### Authors’ contributions

Conceptualization, D.P.; methodology, D.P., S.B., C.P.; investigation, S.B., C.D., C.P., L.M.; writing—original draft, S.B.; review & editing, D.P., S.B., C.P., C.D., L.M.; funding acquisition, D.P.; resources, D.P.; supervision, D.P.

## Acknowledgments

Not applicable

## Figure Captions

Figure 1. The wavelength-dependent phototactic response of *H. symbiolongicarpus* planulae. (**A**) We measured phototaxis in day 3 (72hpf) *H. symbiolongicarpus* planulae using heatless LEDs of known wavelengths in behavioral arenas kept in darkness. Planulae were placed in petri dishes above the LEDs and spatially randomized when ‘before’ photographs were taken. Light of a specific wavelength was then applied for eight hours and an “after” photograph was taken. Images from one replicate are shown. (**B**) For each light condition, we compared the number of respondent (phototactic) larvae to the number of non-respondent larvae using the Welch two-sample t-test. Larvae of *H. symbiolongicarpus* are strongly sensitive to green and blue light, but no significant difference was observed between the number of respondent and non-respondent planulae in red trials. At least 5 separate trials were conducted for each light condition. ** *p* ≤ 0.01; *** *p* ≤ 0.001

Figure 2. Differential gene expression during planula development. (**A**) Multidimensional scaling (MDS) of 12 RNAseq replicates indicates clustering by developmental day. Red = day 1 (24hpf), yellow = day 2 (48hpf), blue = day 3 (72hpf), and green = day 4 (96hpf). (**B**) Pairwise comparison of the number of differentially expressed (DE) transcripts for each developmental day. Most DE transcripts occur between days 1 and 3, and days 1 and 4, while the lowest number of DE transcripts occur between days 2 and 3.

Figure 3. Differential gene expression over development in the *sensory perception of light stimulus* gene set. (**A**) Differentially expressed genes over four days of development examining the *sensory perception of light stimulus* gene set derived from the GSEA (33). Each row is a single transcript labeled by the human gene symbols that are present in the same orthogroup as the planula transcript. High expression is yellow and low is light blue. Grey panels contain abbreviated descriptions of gene functions from UniProt (116). (**B-D**) STRING networks showing significantly differentially expressed genes across four days of development where (**B**) contains all DE genes from all four days, and (**C**) contains subnetworks of DE genes by day. STRING is a database of known and predicted protein-protein interactions (36). In each interaction map, line thickness indicates the strength of data support. Gene symbols are shaded by biological processes (Gene Ontology) or Annotated Keywords (UniProt). (**D**) Node and edge information for each network that includes the PPI enrichment p-values. PPI enrichment p-values double in days 3 and 4, consistent with the expression of phototaxis behavior.

Figure 4. Differential gene expression over development in the *sensory system development gene set*. (**A**) Differentially expressed genes over four days of development examining the *sensory system development* gene set derived from the GSEA (33). Each row is a single transcript labeled by the human gene symbols that are present in the same orthogroup as the planula transcript. High expression is yellow and low is light blue. Grey panels contain abbreviated descriptions of gene functions from UniProt (116). (**B-D**) STRING networks of significantly differentially expressed genes across four days of development where (**B**) contains all DE genes from all four days, and (**C**) contains subnetworks of DE genes by day. In each interaction map, line thickness indicates the strength of data support. Gene symbols are shaded by biological processes (Gene Ontology) or Annotated Keywords (UniProt). (**D**) Node and edge information for each network that includes the PPI enrichment p-values. In contrast to analyses based on the *sensory perception of light stimulus* gene set above, PPI enrichment p-values for the *sensory system development gene set* are highest in day 1, indicating greater functional interactions among genes expressed early in development.

Figure 5. Immunohistochemistry of the nervous system and FMRFamide expression in an *H. symbiolongicarpus* planula larvae. Immunohistochemistry staining of a Day 3 larva (72hpf) where red staining corresponds to acetylated alpha-tubulin of neural cells, magenta corresponds to RFamide, a neurotransmitter involved in relaying photosensory information, green corresponds to F-actin in contractile muscle, and blue corresponds to DAPI staining of nuclei. (**A**) Depicts the whole larva with different merges of the four channels. (**B**) Depicts the zoomed-in view of the aboral plexus which is the site of sensory integration and settlement.

Figure 6. Cnidopsin expressing bipolar neurons comprise the aboral neural plexus in *H. symbiolongicarpus* planula. (**A-D**) Day 3 (72hpf) larva labeled by RNA fluorescent *in situ* hybridization of cnidopsin (green; Hs_t.88569) combined with immunohistochemical staining of neural cells with anti-acetylated alpha-tubulin (red). Nuclear staining is by DAPI (blue). Box in **(A)** depicts section shown in panel (**B**) of the larval *H. symbiolongicarpus* neural plexus where opsin localizes to ciliary regions of bipolar sensory cells, which connect with ganglion neurons to form a plexus. Scale bars = 10um.

Figure 7. Colocalization of photosensory transcripts in the aboral region of the planula. Multiplexed RNA Fluorescent *in situ* hybridization (FISH) of photosensory transcripts including: cnidopsin (green, Hs_t.88569), CNG (red, Hs_t.35573), CRX (magenta, Hs_t.47300), and DAPI (Blue). (**A**) Merged Z-Stack of all channels. (**B**) Each combination of individual transcript expression in single optical slice of a whole mount specimen. (**C-I**) Higher magnification of the aboral region with a focus on the bipolar sensory neurons of the aboral neural plexus. Cnidopsin, CNG ion channels, and CRX strongly localize to the ciliated region of bipolar sensory neurons.

## References

1. Randel N, Jékely G. Phototaxis and the origin of visual eyes. Philos Trans R Soc B Biol Sci. 2016;371(1685).

2. Dunn CW, Giribet G, Edgecombe GD, Hejnol A. Animal phylogeny and its evolutionary implications. Annu Rev Ecol Evol Syst. 2014;45:371–95.

3. Plickert G, Kroiher M, Munck A. Cell proliferation and early differentiation during embryonic development and metamorphosis of Hydractinia echinata. Development. 1988;103(4):795–803.

4. Seipp S, Schmich J, Will B, Schetter E, Plickert G, Leitz T. Neuronal cell death during metamorphosis of Hydractina echinata (Cnidaria, Hydrozoa). Invertebr Neurosci. 2010;10(2):77–91.

5. Weis VM, Keene DR, Buss LW. Biology of Hydractiniid Hydroids. 4. Ultrastructure of the Planula of Hydractinia Echinata. Biol Bull. 1985;168(3):403–18.

6. Plickert G, Schneider B. Neuropeptides and photic behavior in Cnidaria. Hydrobiologia. 2004;530–531:49–57.

7. Katsukura Y, Ando H, David CN, Grimmelikhuijzen CJP, Sugiyama T. Control of planula migration by LWamide and RFamide neuropeptides in Hydractinia echinata. J Exp Biol. 2004;207(11):1803–10.

8. Mason B, Schmale M, Gibbs P, Miller MW, Wang Q, Levay K, et al. Evidence for Multiple Phototransduction Pathways in a Reef-Building Coral. PLoS One. 2012;7(12):1–9.

9. Mason BM, Koyanagi M, Sugihara T, Iwasaki M, Slepak V, Miller DJ, et al. Multiple opsins in a reef-building coral, Acropora millepora. Sci Rep [Internet]. 2023;13(1):1–8. Available from: 10.1038/s41598-023-28476-5

10. Strader ME, Davies SW, Matz M V. Differential responses of coral larvae to the colour of ambient light guide them to suitable settlement microhabitat. R Soc Open Sci. 2015;2(10).

11. Ramirez MD, Pairett AN, Pankey MS, Serb JM, Speiser DI, Swafford AJ, et al. The last common ancestor of most bilaterian animals possessed at least nine opsins. Genome Biol Evol. 2016;8(12):3640–52.

12. Gornik SG, Bergheim BG, Morel B, Stamatakis A, Foulkes NS, Guse A. Photoreceptor Diversification Accompanies the Evolution of Anthozoa. Mol Biol Evol. 2021;38(5):1744– 60.

13. Vöcking O, Kourtesis I, Tumu SC, Hausen H. Co-expression of xenopsin and rhabdomeric opsin in photoreceptors bearing microvilli and cilia. Elife. 2017;6:1–26.

14. Picciani N, Roberts NG, Daly M, Swafford AJM, Kerlin JR, Ramirez MD, et al. Prolific Origination of Eyes in Cnidaria with Co-option of Non-visual Opsins. Curr Biol [Internet]. 2018;28(15):2413–2419.e4. Available from: 10.1016/j.cub.2018.05.055

15. Plachetzki DC, Degnan BM, Oakley TH. The origins of novel protein interactions during animal opsin evolution. PLoS One. 2007;2(10).

16. Plachetzki DC, Fong CR, Oakley TH. Cnidocyte discharge is regulated by light and opsin-mediated phototransduction. BMC Biol. 2012;10(March).

17. Gilbert E, Craggs J, Modepalli V. Gene regulatory network that shaped the evolution of larval apical organ in Cnidaria. Mol Biol Evol [Internet]. 2023;41(1):1–17. Available from: 10.1093/molbev/msad285

18. Sinigaglia C, Busengdal H, Lerner A, Oliveri P, Rentzsch F. Molecular characterization of the apical organ of the anthozoan Nematostella vectensis. Dev Biol [Internet]. 2015;398(1):120–33. Available from: 10.1016/j.ydbio.2014.11.019

19. Gilbert E, Teeling C, Lebedeva T, Pedersen S, Chrismas N, Genikhovich G, et al. Molecular and cellular architecture of the larval sensory organ in the cnidarian Nematostella vectensis. Development. 2022;149(16).

20. Rentzsch F, Fritzenwanker JH, Scholz CB, Technau U. FGF signalling controls formation of the apical sensory organ in the cnidarian Nematostella vectensis. Development. 2008;135(10):1761–9.

21. Marlow H, Tosches MA, Steinmetz PR, Lauri A, Larsson T, Marlow H, et al. Larval body patterning and apical organs are conserved in animal evolution. BMC Biol [Internet]. 2014;12(1):7. Available from: http://www.ncbi.nlm.nih.gov/pubmed/24476105

22. Frank U, Nicotra ML, Schnitzler CE. The colonial cnidarian Hydractinia. Evodevo [Internet]. 2020;11(1):7–12. Available from: 10.1186/s13227-020-00151-0

23. Birch S, Plachetzki D. Multisensory integration by polymodal sensory neurons dictates larval settlement in a brainless cnidarian larva. Mol Ecol. 2023;32(14):3892–907.

24. MacManes MD. The Oyster River Protocol: a multi-assembler and kmer approach for de novo transcriptome assembly. PeerJ. 2018;6:e5428.

25. Patro R, Duggal G, Love MI, Irizarry RA, Kingsford C. Salmon: fast and bias-aware quantification of transcript expression using dual-phase inference. Nat Methods. 2017;14(4):417–9.

26. Chen Y, McCarthy D, Ritchie M, Robinson M, Smyth G, Hall E. edgeR: differential analysis of sequence read count data User’s Guide. R Packag. 2020;(June):1–121.

27. Haas BJ, Papanicolaou A, Yassour M, Grabherr M, Philip D, Bowden J, et al. De novo transcript sequence recostruction from RNA-Seq: reference generation and analysis with Trinity. Vol. 8, Nature protocols. 2013. 1–43 p.

28. Li W, Godzik A. Cd-hit: A fast program for clustering and comparing large sets of protein or nucleotide sequences. Bioinformatics. 2006;22(13):1658–9.

29. Emms DM, Kelly S. OrthoFinder: Phylogenetic orthology inference for comparative genomics. Genome Biol. 2019;20(1):1–14.

30. Sergey N, Sergey K, Arang R, Mikko R, Andrey VB, Alla M, et al. The complete sequence of a human genome. Science (80-). 2022;376(6588):44–53.

31. Kim BY, Wang JR, Miller DE, Barmina O, Delaney E, Thompson A, et al. Highly contiguous assemblies of 101 drosophilid genomes. Elife. 2021;10:1–33.

32. Putnam NH, Srivastava M, Hellsten U, Dirks B, Chapman J, Salamov A, et al. Sea anemone genome reveals ancestral eumetazoan gene repertoire and genomic organization. Science (80-). 2007;317(5834):86–94.

33. Subramanian A, Tamayo P, Mootha VK, Mukherjee S, Ebert BL, Gillette MA, et al. Gene set enrichment analysis: A knowledge-based approach for interpreting genome-wide expression profiles. Proc Natl Acad Sci U S A. 2005;102(43):15545–50.

34. Liberzon A, Birger C, Thorvaldsdóttir H, Ghandi M, Mesirov JP, Tamayo P. The Molecular Signatures Database Hallmark Gene Set Collection. Cell Syst. 2015;1(6):417–25.

35. Liberzon A, Subramanian A, Pinchback R, Thorvaldsdóttir H, Tamayo P, Mesirov JP. Molecular signatures database (MSigDB) 3.0. Bioinformatics. 2011;27(12):1739–40.

36. Szklarczyk D, Gable AL, Nastou KC, Lyon D, Kirsch R, Pyysalo S, et al. The STRING database in 2021: Customizable protein-protein networks, and functional characterization of user-uploaded gene/measurement sets. Nucleic Acids Res. 2021;49(D1):D605–12.

37. Müller WA, Leitz T. Metamorphosis in the Cnidaria. Can J Zool [Internet]. 2002;80(10):1755–71. Available from: 10.1139/z02-130

38. Koonin E V. Orthologs, Paralogs, and Evolutionary Genomics. Annu Rev Genet. 2005;39(1):309–38.

39. Koyanagi M, Takano K, Tsukamoto H, Ohtsu K, Tokunaga F, Terakita A. Jellyfish vision starts with cAMP signaling mediated by opsin-Gs cascade. Proc Natl Acad Sci [Internet]. 2008;105(40):15576–80. Available from: 10.1073/pnas.0806215105

40. Plachetzki DC, Fong CR, Oakley TH. The evolution of phototransduction from an ancestral cyclic nucleotide gated pathway. Proc R Soc B Biol Sci. 2010;277(1690):1963–9.

41. Mason BM, Cohen JH. Long-Wavelength Photosensitivity in Coral planula larvae. Biol Bull. 2012;222(2):88–92.

42. McCulloch KJ, Babonis LS, Liu A, Daly CM, Martindale MQ, Koenig KM. Nematostella vectensis exemplifies the exceptional expansion and diversity of opsins in the eyeless Hexacorallia. Evodevo [Internet]. 2023;14(1):1–19. Available from: 10.1186/s13227-023-00218-8

43. Quiroga Artigas G, Lapébie P, Leclère L, Takeda N, Deguchi R, Jékely G, et al. A gonad-expressed opsin mediates light-induced spawning in the jellyfish Clytia. Elife. 2018;7:1– 22.

44. Leach WB, Reitzel AM. Decoupling behavioral and transcriptional responses to color in an eyeless cnidarian. BMC Genomics. 2020;21(1):1–15.

45. Szklarczyk D, Gable AL, Lyon D, Junge A, Wyder S, Huerta-Cepas J, et al. STRING v11: Protein-protein association networks with increased coverage, supporting functional discovery in genome-wide experimental datasets. Nucleic Acids Res. 2019;47(D1):D607– 13.

46. Hao H, Kim DS, Klocke B, Johnson KR, Cui K, Gotoh N, et al. Transcriptional regulation of rod photoreceptor homeostasis revealed by in vivo NRL targetome analysis. PLoS Genet. 2012;8(4).

47. Hennig AK, Peng G-H, Chen S. Regulation of photoreceptor gene expression by Crx-associated transcription factor network. Brain Res. 2008;1192:114–33.

48. Henrique D, Hirsinger E, Adam J, Roux I Le, Pourquié O, Ish-Horowicz D, et al. Maintenance of neuroepithelial progenitor cells by Delta-Notch signalling in the embryonic chick retina. Curr Biol. 1997;7(9):661–70.

49. Martin VJ. Reorganization of the nervous system during metamorphosis of a hydrozoan planula. Invertebr Biol. 2000;119(3):243–53.

50. Schwoerer-Böhning B, Kroiher M, Müller WA. Signal transmission and covert prepattern in the metamorphosis of Hydractinia echinata (Hydrozoa). Roux’s Arch Dev Biol. 1990;198(5):245–51.

51. Yau KW, Hardie RC. Phototransduction Motifs and Variations. Cell. 2009;139(2):246–64.

52. Brzezinski JA, Reh TA. Photoreceptor cell fate specification in vertebrates. Dev. 2015;142(19):3263–73.

53. Birch S, Picciani N, Oakley TH, Plachetzki D. Cnidarians: Diversity and Evolution of Cnidarian Visual Systems. In: Buschbeck E, Bok M, editors. Distributed Vision: From Simple Sensor to Sophisticated Combination Eyes. Springer; 2023. p. 21–47.

54. Brodrick E, Jékely G. Photobehaviours guided by simple photoreceptor systems. Anim Cogn [Internet]. 2023;(0123456789). Available from: 10.1007/s10071-023-01818-6

55. Forward RB. Diel Vertical Migration: Zooplankton Photobiology and Behaviour. In: Oceanography and Marine Biology, An Annual Review. 1988. p. 361–93.

56. Thorson G. Light as an ecological factor in the dispersal and settlement of larvae of marine bottom invertebrates. Ophelia. 1964;1(1):167–208.

57. Mundy CN, Babcock RC. Role of light intensity and spectral quality in coral settlement: Implications for depth-dependent settlement? J Exp Mar Bio Ecol. 1998;223(2):235–55.

58. Guertin S, Kass-Simon G. Extraocular spectral photosensitivity in the tentacles of Hydra vulgaris. Comp Biochem Physiol-Part A Mol Integr Physiol [Internet]. 2015;184:163–70. Available from: 10.1016/j.cbpa.2015.02.016

59. Schijfsma K. Observations on Hydractinia Echinata (Flem. and Eupagurus Bernhardus(L.). Arch Neerl Zool. 1934;1(1):261–314.

60. Hodin J, Ferner MC, Heyland A, Gaylord B. I feel that! fluid dynamics and sensory aspects of larval settlement across scales. Evolutionary Ecology of Marine Invertebrate Larvae. 2018. 190–207 p.

61. Kingsford MJ, Leis JM, Shanks A, Lindeman KC, Morgan SG, Pineda J. Sensory environments, larval abilities and local self-recruitment. Bull Mar Sci. 2002;70(1 SUPPL.):309–40.

62. Say TE, Degnan SM. Molecular and behavioural evidence that interdependent photo-and chemo-sensory systems regulate larval settlement in a marine sponge. Mol Ecol. 2019;(29):247–61.

63. Technau U, Steele RE. Evolutionary crossroads in developmental biology: Cnidaria. Development. 2012;139(23):4491–4491.

64. Bosch TCG, Klimovich A, Domazet-Lošo T, Gründer S, Holstein TW, Jékely G, et al. Back to the Basics: Cnidarians Start to Fire. Trends Neurosci. 2017;40(2):92–105.

65. Taranova O V., Magness ST, Fagan BM, Wu Y, Surzenko N, Hutton SR, et al. SOX2 is a dose-dependent regulator of retinal neural progenitor competence. Genes Dev. 2006;20(9):1187–202.

66. Chrysostomou E, Flici H, Gornik SG, Salinas-Saavedra M, Gahan JM, McMahon ET, et al. A cellular and molecular analysis of SoxB-driven neurogenesis in a cnidarian. Elife. 2022;11:1–24.

67. Richards GS, Rentzsch F. Regulation of Nematostella neural progenitors by SoxB, Notch and bHLH genes. Dev. 2015;142(19):3332–42.

68. Javed A, Mattar P, Lu S, Kruczek K, Kloc M, Gonzalez-Cordero A, et al. Pou2f1 and Pou2f2 cooperate to control the timing of cone photoreceptor production in the developing mouse retina. Dev. 2020;147(18).

69. Gold DA, Gates RD, Jacobs DK. The early expansion and evolutionary dynamics of POU class genes. Mol Biol Evol. 2014;31(12):3136–47.

70. Cathriona Millane R, Kanska J, Duffy DJ, Seoighe C, Cunningham S, Plickert G, et al. Induced stem cell neoplasia in a cnidarian by ectopic expression of a POU domain transcription factor. Development. 2011;138(12):2429–39.

71. Ozment E, Tamvacakis AN, Zhou J, Rosiles-Loeza PY, Escobar-Hernandez EE, Fernandez-Valverde SL, et al. Cnidarian hair cell development illuminates an ancient role for the class IV POU transcription factor in defining mechanoreceptor identity. Elife. 2021;10:1– 34.

72. Peng GH, Ahmad O, Ahmad F, Liu J, Chen S. The photoreceptor-specific nuclear receptor Nr2e3 interacts with Crx and exerts opposing effects on the transcription of rod versus cone genes. Hum Mol Genet. 2005;14(6):747–64.

73. Roduit R, Escher P, Schorderet DF. Mutations in the DNA-binding domain of NR2E3 affect in vivo dimerization and interaction with CRX. PLoS One. 2009;4(10):1–12.

74. Siemens J, Lillo C, Dumont RA, Reynolds A, Williams DS, Gillespie PG, et al. Cadherin 23 Is a component of the tip link in hair-cell stereocilla. Nature. 2004;428(6986):950–5.

75. Liu X, Bulgakov O V., Darrow KN, Pawlyk B, Adamian M, Liberman MC, et al. Usherin is required for maintenance of retinal photoreceptors and normal development of cochlear hair cells. Proc Natl Acad Sci U S A. 2007;104(11):4413–8.

76. Dona M, Slijkerman R, Lerner K, Broekman S, Wegner J, Howat T, et al. Usherin defects lead to early-onset retinal dysfunction in zebrafish. Exp Eye Res. 2018;173(May):148–59.

77. Adato A, Lefèvre G, Delprat B, Michel V, Michalski N, Chardenoux S, et al. Usherin, the defective protein in Usher syndrome type IIA, is likely to be a component of interstereocilia ankle links in the inner ear sensory cells. Hum Mol Genet. 2005;14(24):3921–32.

78. Tang PC, Watson GM. Cadherin-23 may be dynamic in hair bundles of the model sea anemone Nematostella vectensis. PLoS One. 2014;9(1).

79. Bolz H, Brederlow B Von, Ramírez A, Bryda EC, Kutsche K, Gerd H, et al. Mutation of CDH23, encoding a new member of the cadherin gene family, causes Usher syndrome type 1D Hanno. Nat Genet. 2001;27(january).

80. Tucker RP. Expression of usherin in the anthozoan Nematostella vectensis. Biol Bull. 2010;218(2):105–12.

81. Jongbloets BC, Jeroen Pasterkamp R. Semaphorin signalling during development. Dev. 2014;141(17):3292–7.

82. Naeem MA, Chavali VRM, Ali S, Iqbal M, Riazuddin S, Khan SN, et al. GNAT1 associated with autosomal recessive congenital stationary night blindness. Investig Ophthalmol Vis Sci. 2012;53(3):1353–61.

83. McGee JA, Goodyear RJ, McMillan DR, Stauffer EA, Holt JR, Locke KG, et al. The very large G-protein-coupled receptor VLGR1: A component of the ankle link complex required for the normal development of auditory hair bundles. J Neurosci. 2006;26(24):6543–53.

84. Piatigorsky J, Kozmik Z. Cubozoan jellyfish: An evo/devo model for eyes and other sensory systems. Int J Dev Biol. 2004;48(8–9):719–29.

85. Revilla-i-Domingo R, Veedin Rajan VB, Waldherr M, Prohaczka G, Musset H, Orel L, et al. Characterization of cephalic and non-cephalic sensory cell types provides insight into joint photo-and mechanoreceptor evolution. bioRxiv. 2021;1–65.

86. Cosgrove D, Zallocchi M. Usher protein functions in hair cells and photoreceptors. Int J Biochem Cell Biol [Internet]. 2014;46(1):80–9. Available from: 10.1016/j.biocel.2013.11.001

87. Arendt D. The evolution of cell types in animals: Emerging principles from molecular studies. Nat Rev Genet. 2008;9(11):868–82.

88. Sherman PM, Sun H, Macke JP, Williams J, Smallwood PM, Nathans J. Identification and characterization of a conserved family of protein serine/threonine phosphatases homologous to Drosophila retinal degeneration C (rdgC). Proc Natl Acad Sci U S A. 1997;94(21):11639–44.

89. Vinós J, Jalink K, Hardy RW, Britt SG, Zuker CS. A G protein-coupled receptor phosphatase required for rhodopsin function. Science (80-). 1997;277(5326):687–90.

90. Shieh BH. Molecular genetics of retinal degeneration a drosophila perspective. Fly (Austin). 2011;5(4):356–68.

91. Nishiguchi KM, Sandberg MA, Kooijman AC, Martemyanov KA, Pott JWR, Hagstrom SA, et al. Defects in RGS9 or its anchor protein R9AP in patients with slow photoreceptor deactivation. Nature. 2004;427(6969):75–8.

92. McEntaffer RL, Natochin M, Artemyev NO. Modulation of transducin GTPase activity by chimeric RGS16 and RGS9 regulators of G protein signaling and the effector molecule. Biochemistry. 1999;38(16):4931–7.

93. Chen CK, Zhang K, Church-Kopish J, Huang W, Zhang H, Chen YJ, et al. Characterization of human GRK7 as a potential cone opsin kinase. Mol Vis. 2001;7(December):305–13.

94. Shen D, Jiang M, Hao W, Li T, Salazar M, Fong HKW. A Human Opsin-Related Gene That Encodes a Retinaldehyde-Binding Protein. Biochemistry. 1994;33(44):13117–25.

95. Garm A, Svaerke JE, Pontieri D, Oakley TH. Expression of Opsins of the Box Jellyfish Tripedalia cystophora Reveals the First Photopigment in Cnidarian Ocelli and Supports the Presence of Photoisomerases. Front Neuroanat. 2022;16(August):1–13.

96. Fei Y, Hughes TE. Transgenic expression of the jellyfish green fluorescent protein in the cone photoreceptors of the mouse. Vis Neurosci. 2001;18(4):615–23.

97. Liegertová M, Pergner J, Kozmiková I, Fabian P, Pombinho AR, Strnad H, et al. Cubozoan genome illuminates functional diversification of opsins and photoreceptor evolution. Sci Rep [Internet]. 2015;5:1–19. Available from: 10.1038/srep11885

98. Cote RH, Gupta R, Irwin MJ, Wang X. Photoreceptor Phosphodiesterase (PDE6): Structure, Regulatory Mechanisms, and Implications for Treatment of Retinal Diseases. Adv Exp Med Biol. 2022;1371:33–59.

99. Kawamura S, Tachibanaki S. Rod and cone photoreceptors: Molecular basis of the difference in their physiology. Comp Biochem Physiol - A Mol Integr Physiol. 2008;150(4):369–77.

100. van Wyk M, Kleinlogel S. A visual opsin from jellyfish enables precise temporal control of G protein signalling. Nat Commun. 2023;14(1).

101. Kozmik Z, Ruzickova J, Jonasova K, Matsumoto Y, Vopalensky P, Kozmikova I, et al. Assembly of the cnidarian camera-type eye from vertebrate-like components. Proc Natl Acad Sci. 2008;105(26):8989–93.

102. Mercando N, Lytle C. Specificity in the association between Hydractinia echinata and sympatric species of hermit crabs. Biol Bull. 1980;159:337–48.

103. Nagel-Wolfrum K, Fadl BR, Becker MM, Wunderlich KA, Schäfer J, Sturm D, et al. Expression and subcellular localization of USH1C/harmonin in human retina provides insights into pathomechanisms and therapy. Hum Mol Genet. 2023;32(3):431–49.

104. García-Añoveros J, Clancy JC, Foo CZ, García-Gómez I, Zhou Y, Homma K, et al. Tbx2 is a master regulator of inner versus outer hair cell differentiation. Nature. 2022;605(7909):298–303.

105. Angueyra JM, Kunze VP, Patak LK, Kim H, Kindt K, Li W. Transcription factors underlying photoreceptor diversity. Elife. 2023;12:1–30.

106. Nakanishi N, Renfer E, Technau U, Rentzsch F. Nervous systems of the sea anemone Nematostella vectensis are generated by ectoderm and endoderm and shaped by distinct mechanisms. Development [Internet]. 2012;139(2):347–57. Available from: 10.1242/dev.071902

107. Nakanishi N, Yuan D, Jacobs DK, Hartenstein V. Early development, pattern, and reorganization of the planula nervous system in Aurelia (Cnidaria, Scyphozoa). Dev Genes Evol. 2008;218(10):511–24.

108. Seipp S, Schmich J, Kehrwald T, Leitz T. Metamorphosis of Hydractinia echinata - Natural versus artificial induction and developmental plasticity. Dev Genes Evol. 2007;217(5):385–94.

109. Martin VJ. Development of Nerve Cells in Hydrozoan Planulae: I. Differentiation of Ganglionic Cells. Biol Bull. 1988;174(3):319–29.

110. Schmich J, Trepel S, Leitz T. The role of GLWamides in metamorphosis of Hydractinia echinata. Dev Genes Evol. 1998;208:267–73.

111. Plickert G. Proportion-altering factor (PAF) stimulates nerve cell formation in Hydractinia echinata. Cell Differ Dev. 1989;26(1):19–27.

112. Gajewski M, Leitz T, Schloßherr J, Plickert G. LWamides from Cnidaria constitute a novel family of neuropeptides with morphogenetic activity. Roux’s Arch Dev Biol. 1996;205(5– 6):232–42.

113. Leitz T. Biochemical and cytological bases of metamorphosis in Hydractinia echinata. Mar Biol Int J Life Ocean Coast Waters. 1993;116(4):559–64.

114. Piraino S, Zega G, Di Benedetto C, Leone A, Dell’Anna A, Pennati R, et al. Complex neural architecture in the diploblastic larva of Clava multicornis (Hydrozoa, Cnidaria). J Comp Neurol. 2011;519(10):1931–51.

115. Shichida Y, Matsuyama T. Evolution of opsins and phototransduction. Philos Trans R Soc B Biol Sci. 2009;364(1531):2881–95.

116. The UniProt Consortium. UniProt: the universal protein knowledgebase in 2021. Nucleic Acids Res. 2021;49(D1):Pages D480–D489.

