## Supplementary figures and images for "Phototactic preference and its genetic basis in the planulae of the colonial Hydrozoan Hydractinia symbiolongicarpus"

### Fig_2.pdf

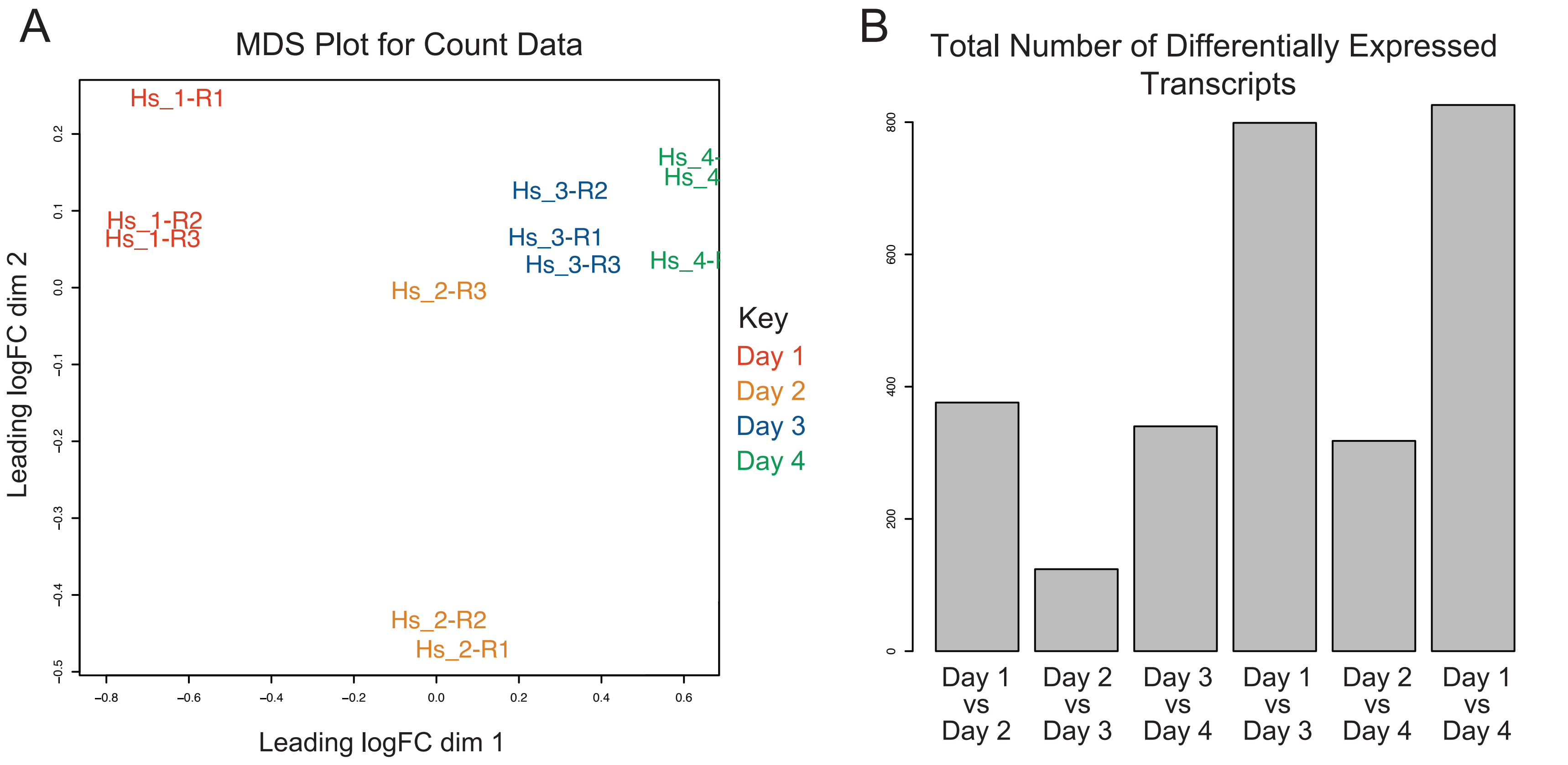

### Fig_4.pdf

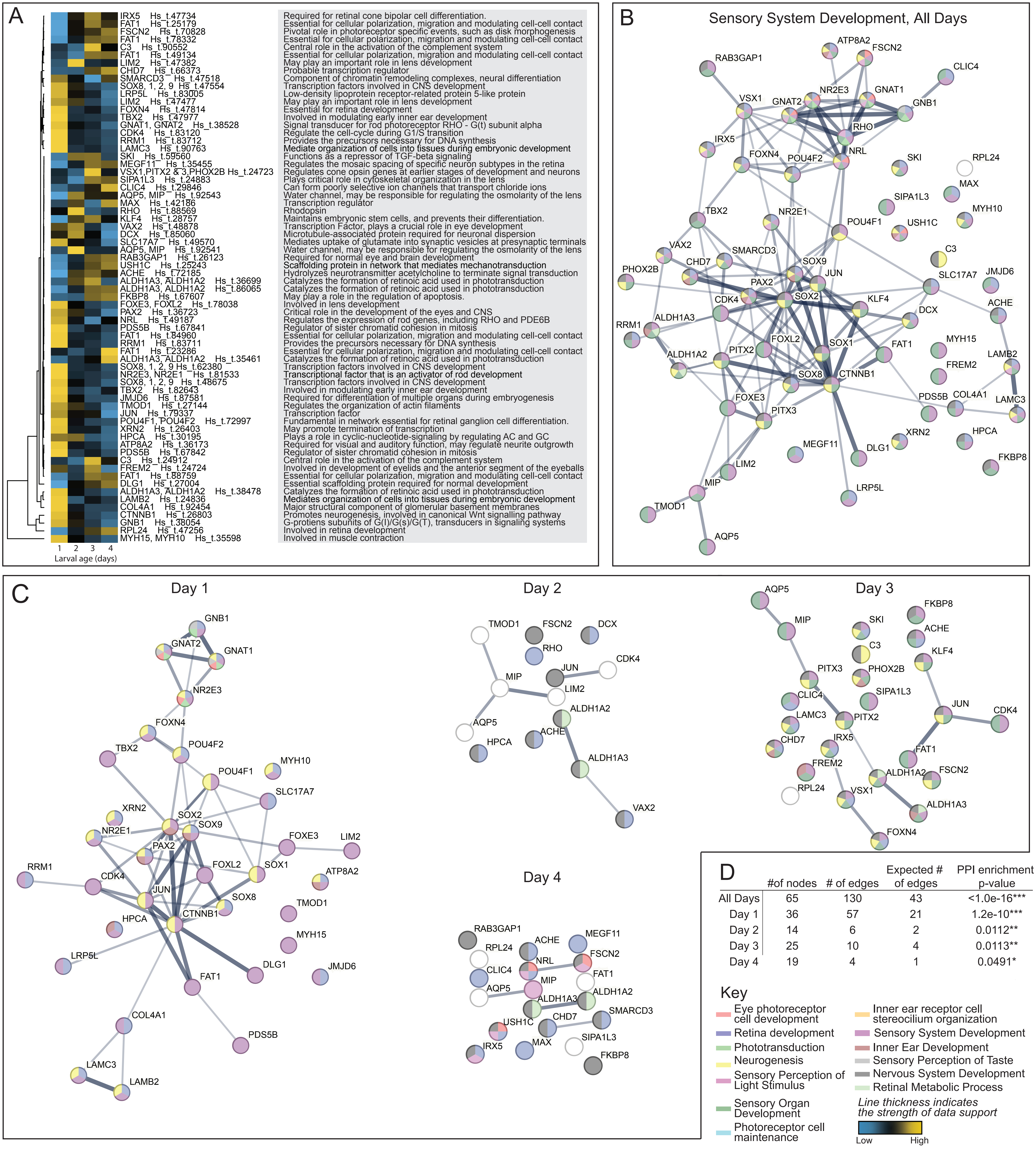

### Fig_5.pdf

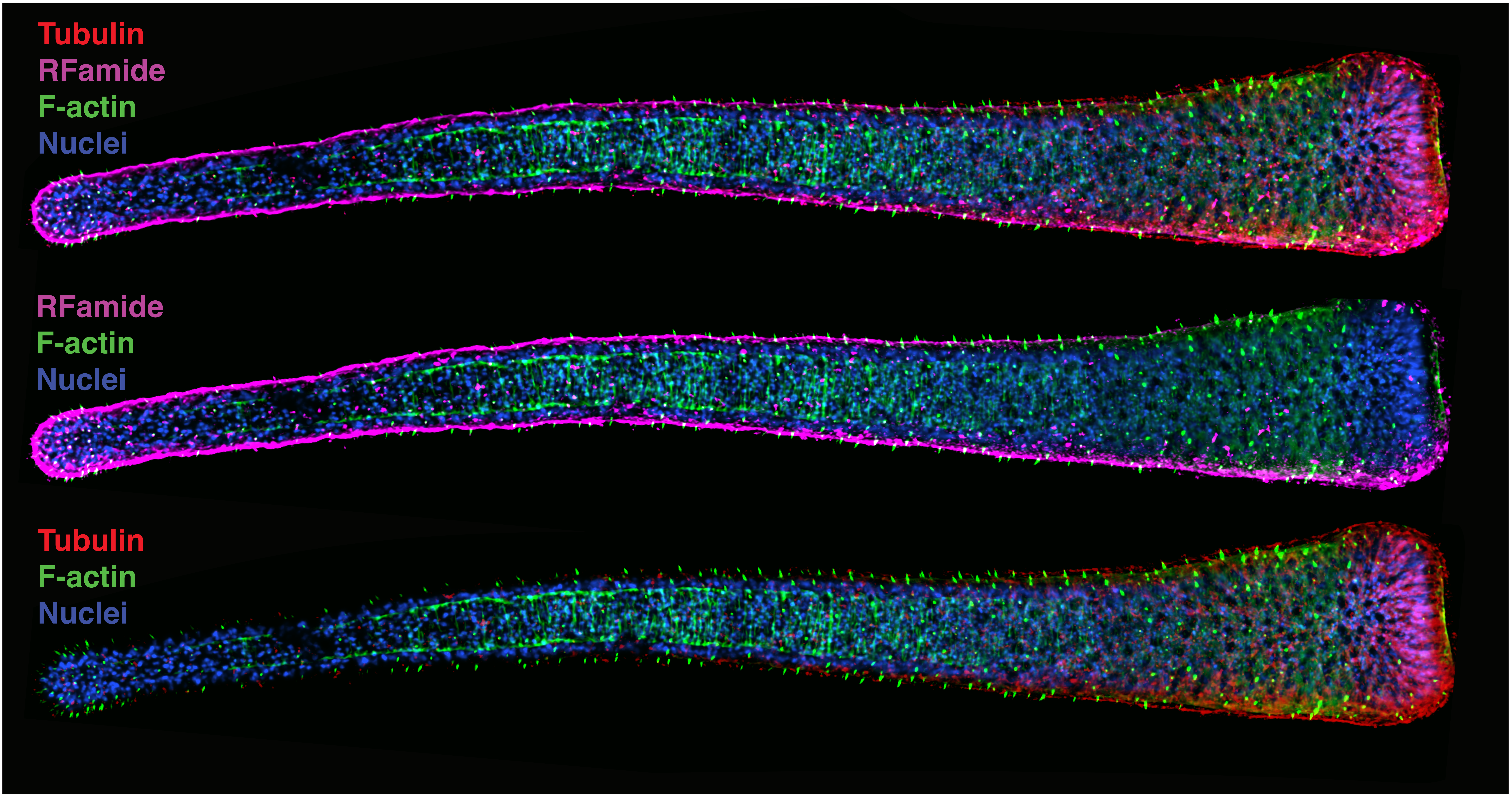

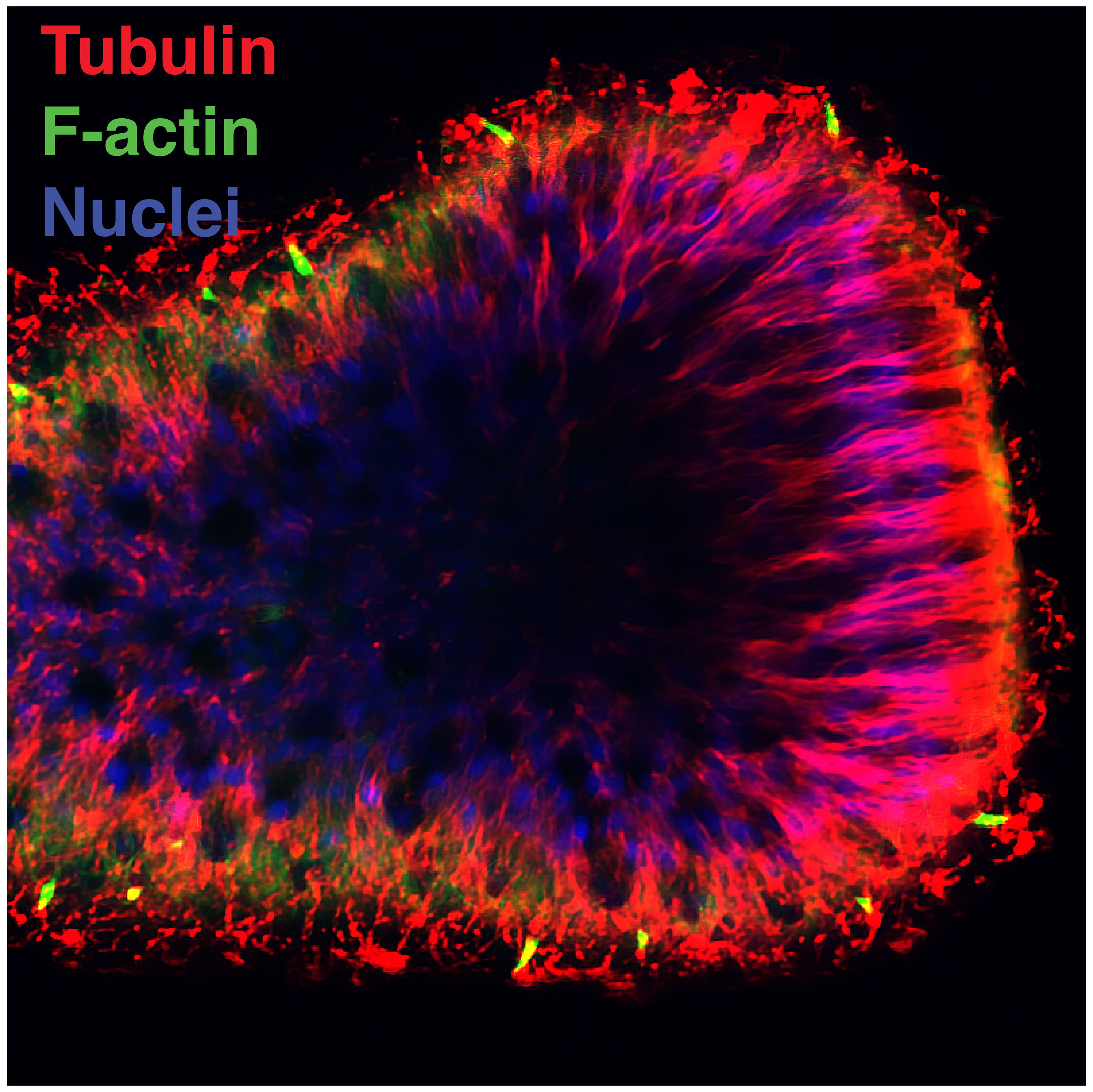

### Fig_6.pdf

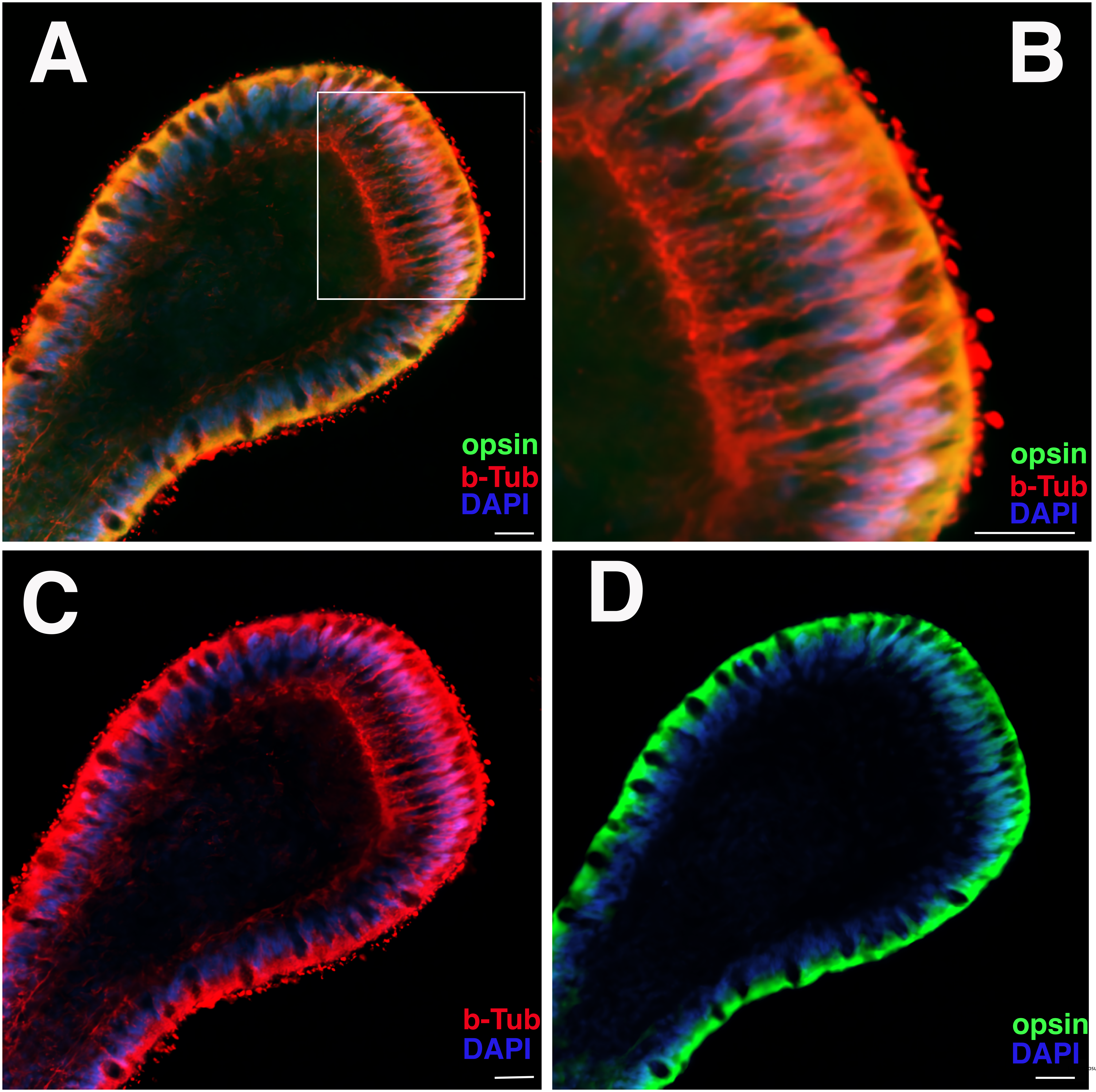

### Fig_7.pdf

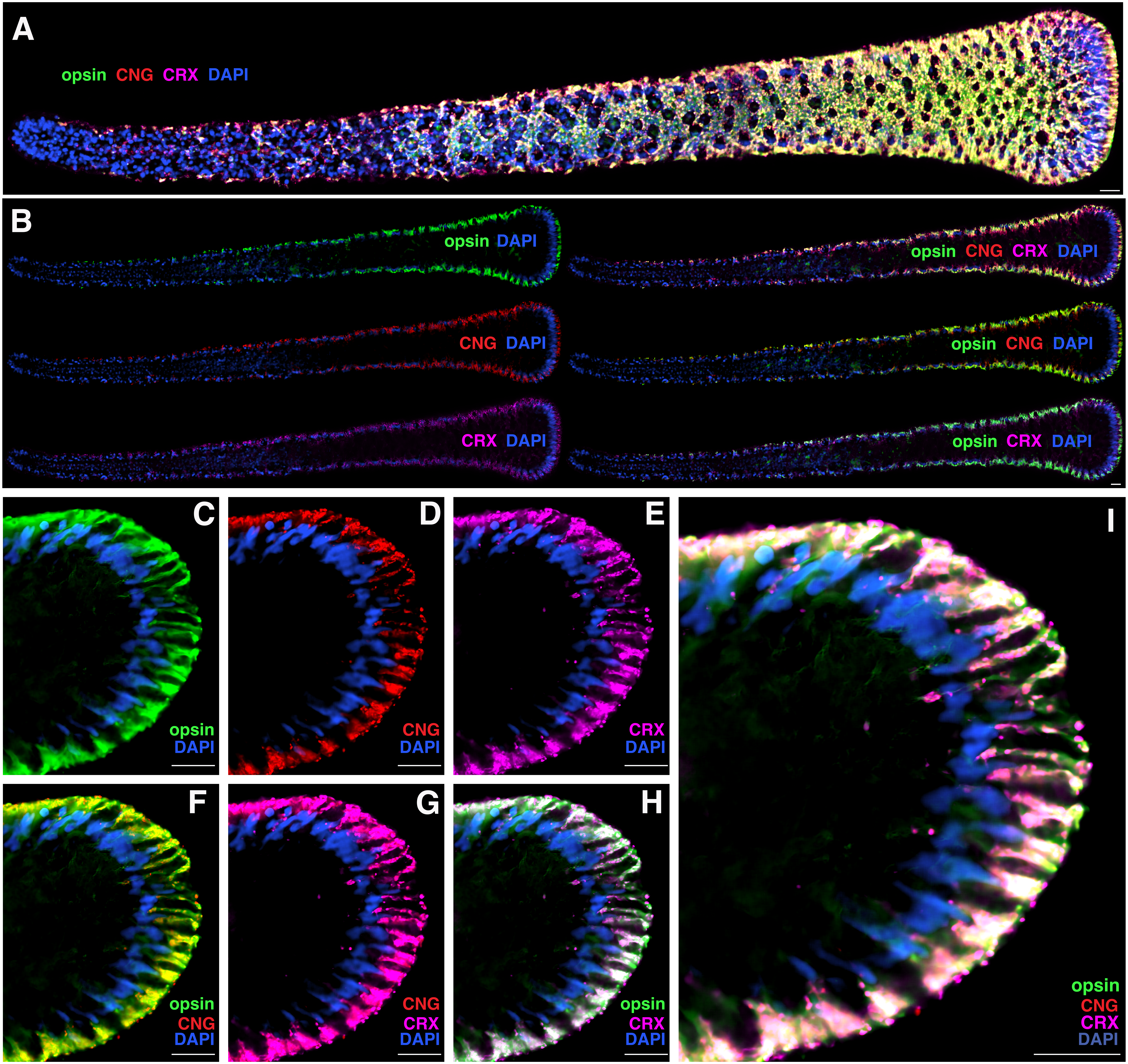

### Supp_Fig_1.pdf

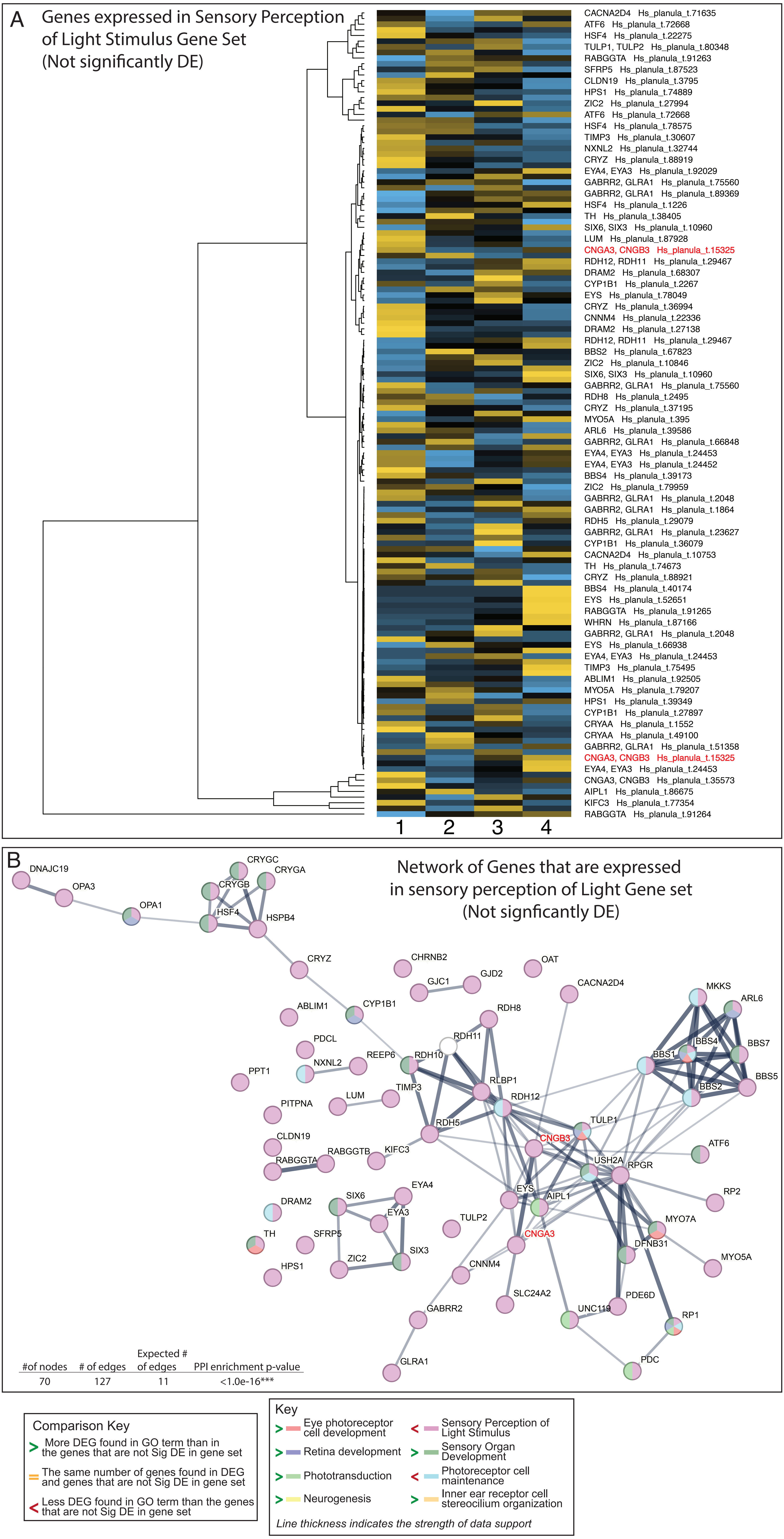

### Supp_Fig_3.pdf

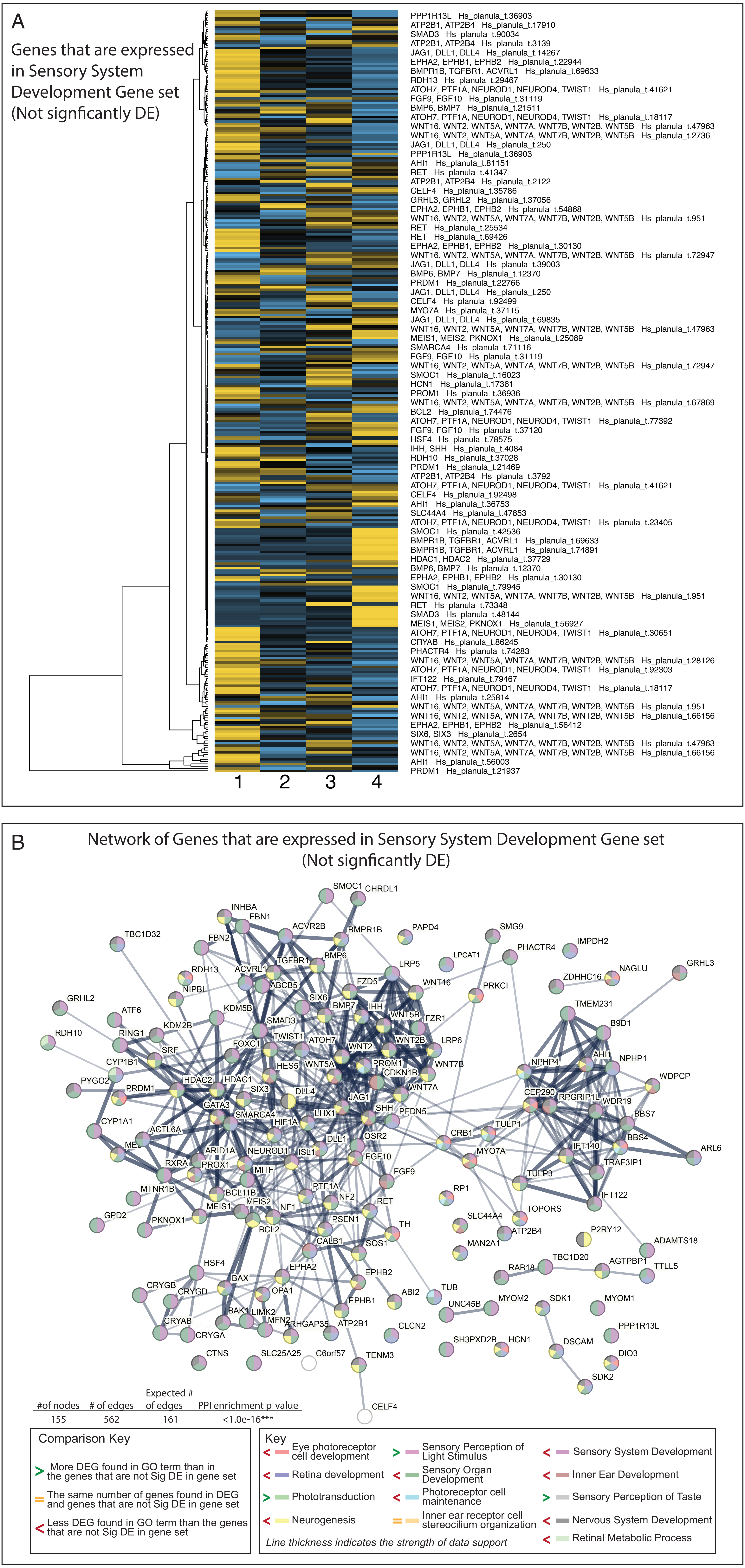

### Supp_Fig_5.pdf

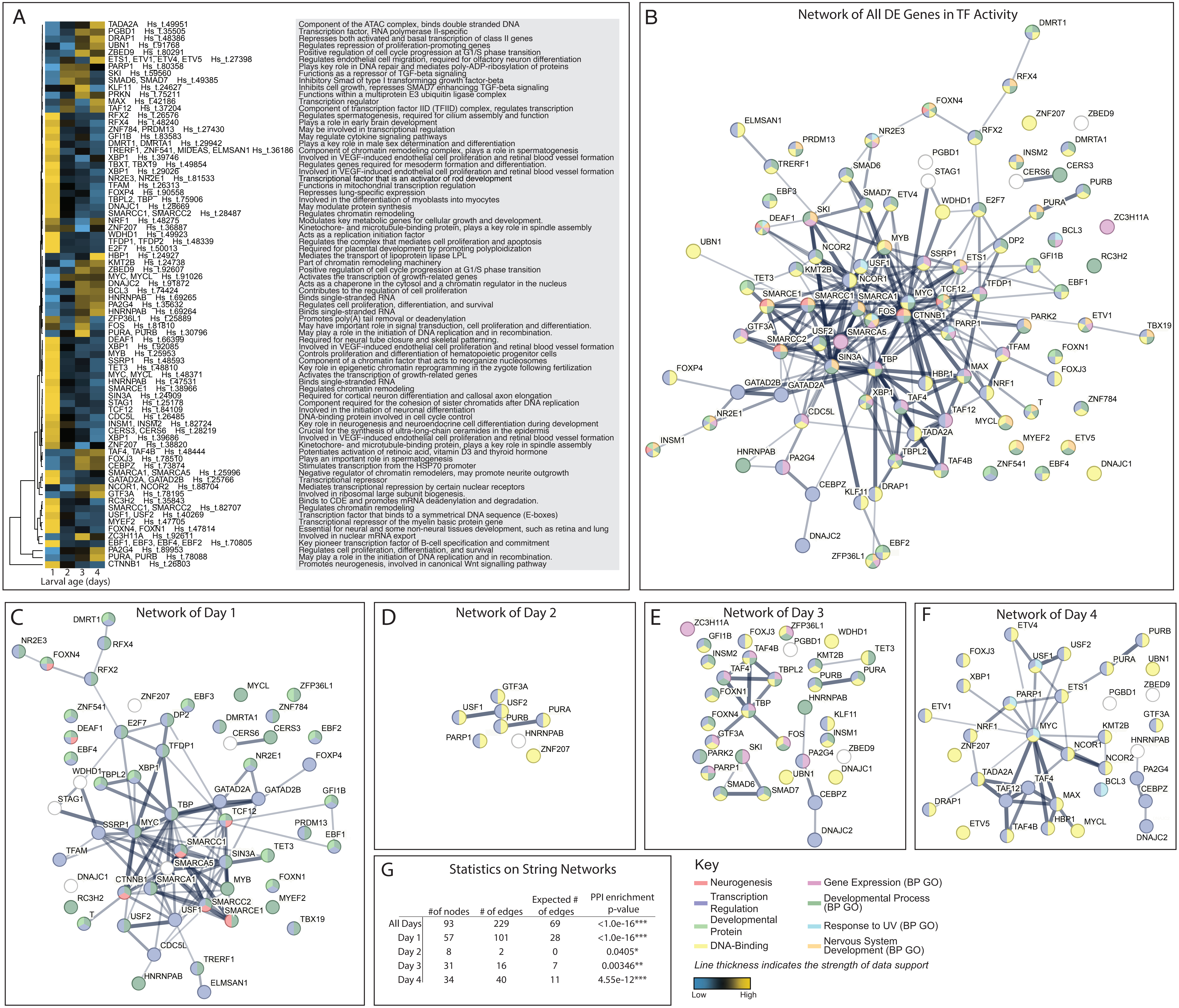


| | #of nodes | # of edges | Expected # of edges | PPI enrichment p-value |
|---|---|---|---|---|
| All Days | 93 | 229 | 69 | <1.0e-16*** |
| Day 1 | 57 | 101 | 28 | <1.0e-16*** |
| Day 2 | 8 | 2 | 0 | 0.0405* |
| Day 3 | 31 | 16 | 7 | 0.00346** |
| Day 4 | 34 | 40 | 11 | 4.55e-12*** |

### Supp_Fig_6.pdf

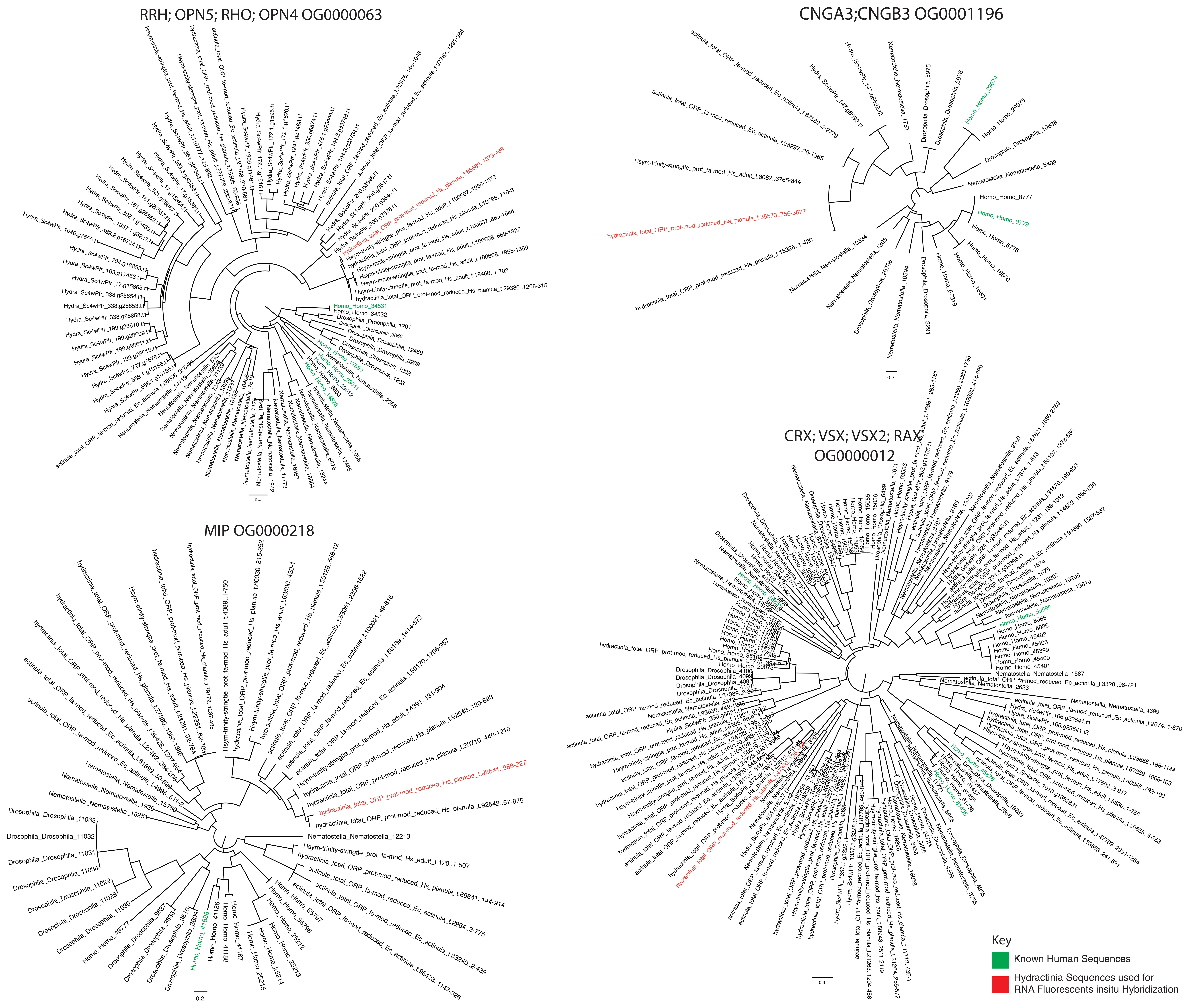
