## Supplementary material for "Phototactic preference and its genetic basis in the planulae of the colonial Hydrozoan Hydractinia symbiolongicarpus": Figure 3: FIg_3.pdf

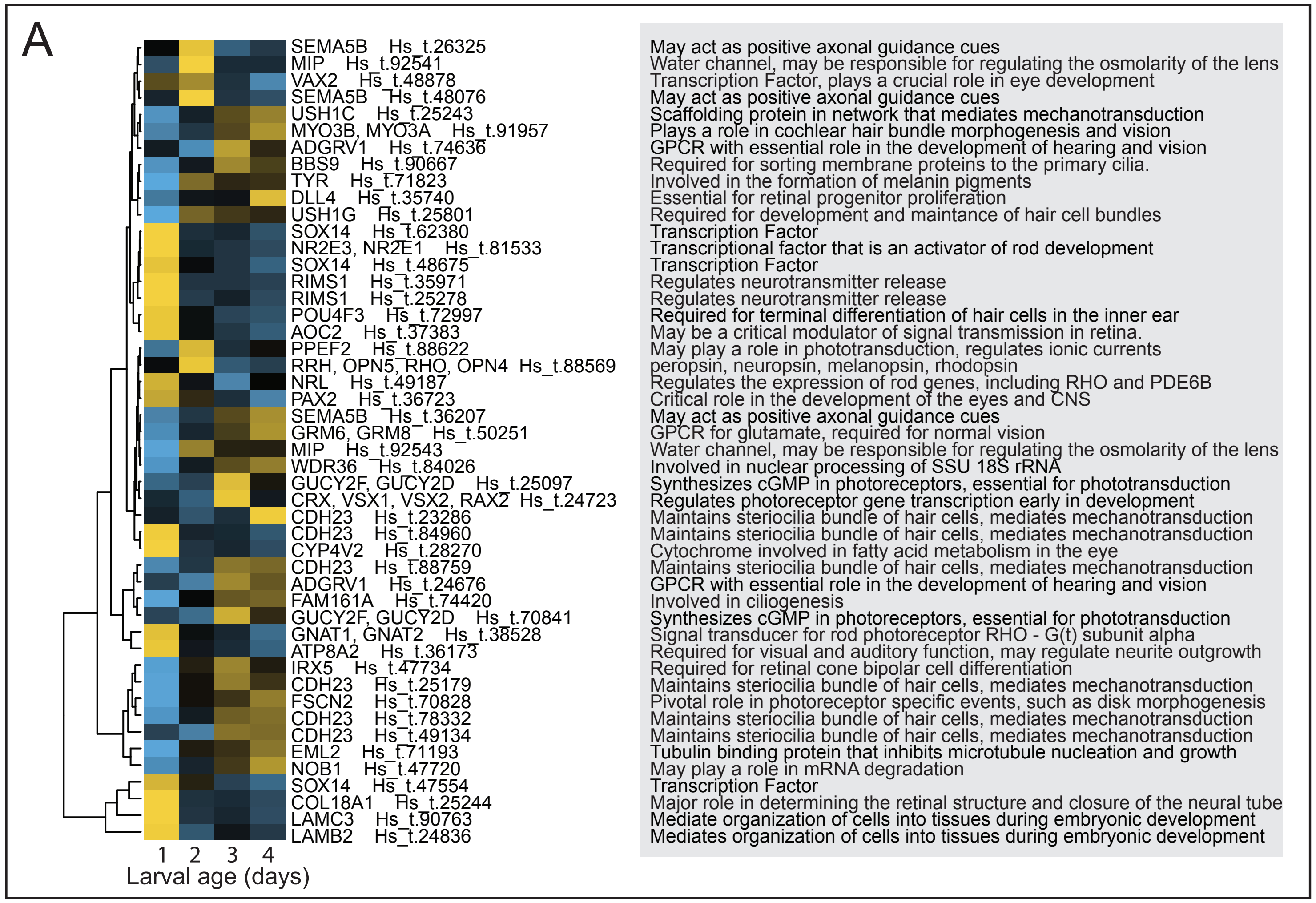


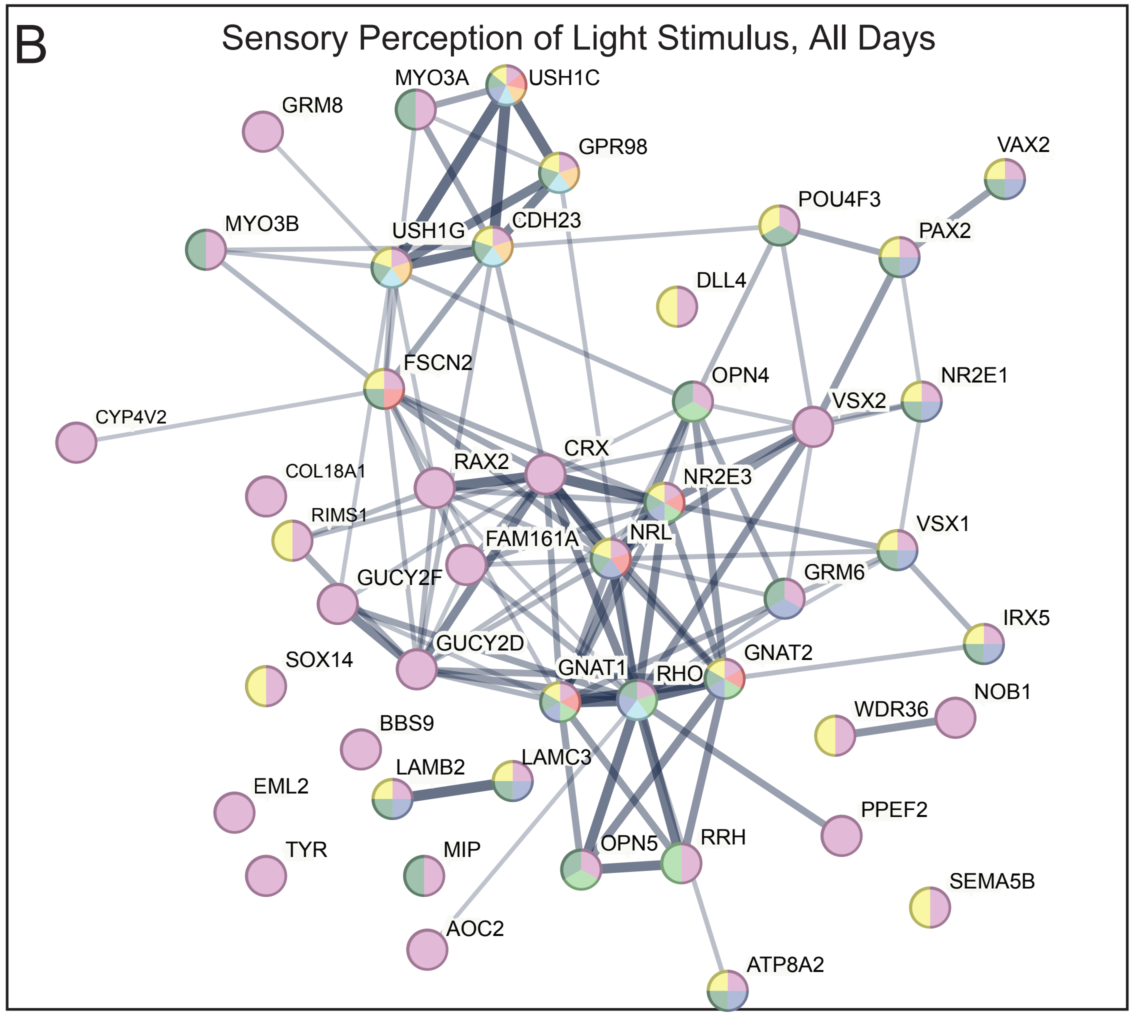


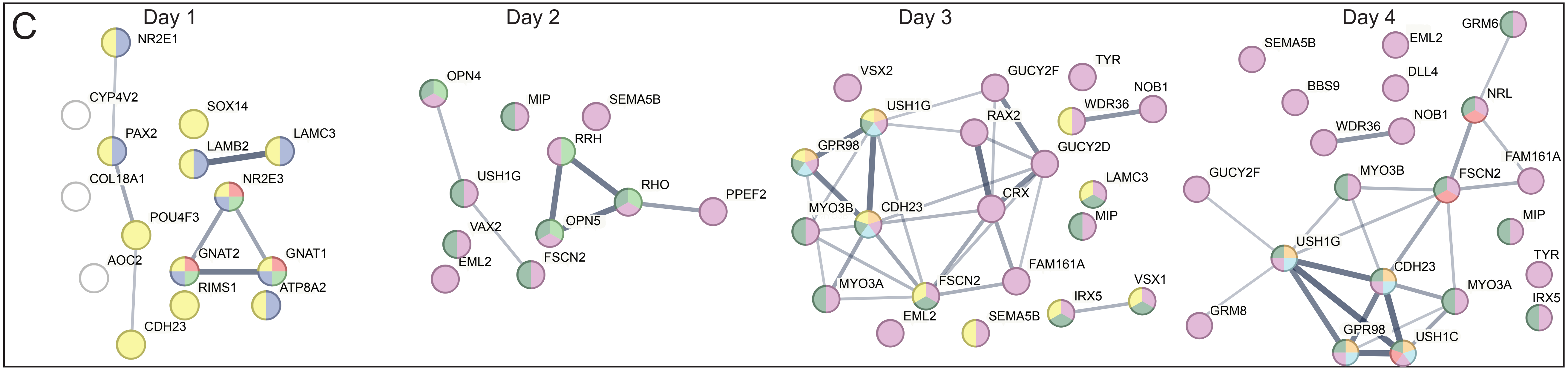


D

### Statistics on String Networks

| | #of nodes | # of edges | Expected # of edges | PPI enrichment p-value |
|---|---|---|---|---|
| All Days | 46 | 109 | 7 | <1.0e-16*** |
| Day 1 | 15 | 7 | 1 | 4.88e-06*** |
| Day 2 | 11 | 6 | 0 | 5.46e-0.8*** |
| Day 3 | 21 | 27 | 2 | <1.0e-16*** |
| Day 4 | 21 | 22 | 2 | <1.0e-16*** |

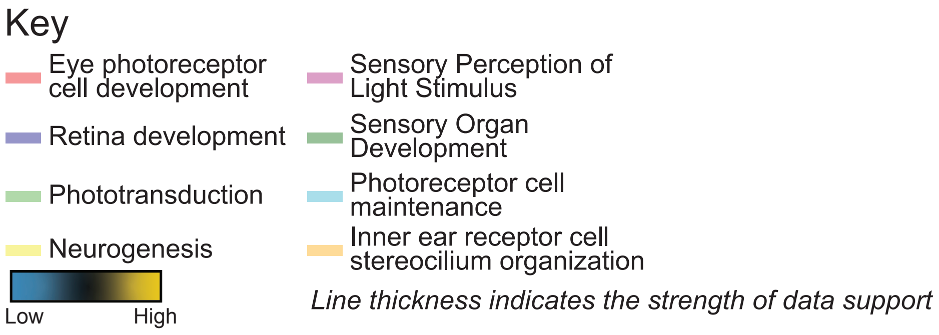
