## Supplementary Information for "Phototactic preference and its genetic basis in the planulae of the colonial Hydrozoan Hydractinia symbiolongicarpus"

### 1.1 Supplementary Figures

**Supplementary Figure 1.** Genes expressed, but not significantly differentially expressed in the *sensory perception of light* gene set. (**A**) Gene expression (not significantly differentially expressed) over four days of development examining the *sensory perception of light stimulus* gene set derived from GSEA (33). Each row is a single transcript labeled by the human gene symbols that are present in the same orthogroup as the planula transcript. High expression = yellow, low expression = light blue. Red text indicates transcripts of CNG found in this dataset, where transcript Hs_t.35573 was used as the target sequence for designing RNA fluorescent in situ hybridization probes. (**B**) STRING network showing expressed genes across four days of development. STRING is a database of known and predicted protein-protein interactions (36). In the interaction map, line thickness indicates the strength of data support. Gene symbols are shaded by biological processes (Gene Ontology) or Annotated Keywords (UniProt). Node and edge information is given for the network and includes the protein-protein interaction (PPI) enrichment p-values. Green > sign indicates that there are more significantly DE genes found in the particular GO term than genes that are expressed (not significantly DE). The yellow = sign means there are the same number of genes that are significantly DE in this GO term as genes that are just expressed (not significantly DE). The red < sign indicates that there are fewer significantly DE genes in this GO term compared to genes that are just expressed (not significantly DE).

**Supplementary Figure 2.** STRING network of genes not expressed in the *sensory perception of light* gene set (36). This STRING network shows the orthogroups of genes labeled by human gene symbols that are in the gene set that did not contain an *H. symbiolongicarpus* transcript. The key contains the color-coding information for each gene in the STRING networks describing selected biological processes (Gene Ontology) or Annotated Keywords (UniProt).

**Supplementary Figure 3.** Genes expressed, but not significantly differentially in the *sensory system development* gene set. (**A**) Gene expression (not significantly DE) over four days of development examining the *sensory system development* gene set derived from the GSEA (33). Each row is a single transcript labeled by the human gene symbols that are present in the same orthogroup as the planula transcript. High expression is yellow and low is light blue. (**B**) STRING network showing expressed genes across four days of development. STRING is a database of known and predicted protein-protein interactions (36). In the interaction map, line thickness indicates the strength of data support. Gene symbols are shaded by biological processes (Gene Ontology) or Annotated Keywords (UniProt). Node and edge information is given for the network and includes the protein-

protein interaction (PPI) enrichment p-values. Green > sign indicates that there are more significantly DE genes found in the particular GO term than genes that are expressed (not significantly DE). The yellow = sign means there are the same number of genes that are significantly DE in this GO term as genes that are just expressed (not significantly DE). The red < sign indicates that there are fewer significantly DE genes in this GO term compared to genes that are just expressed (not significantly DE).

**Supplementary Figure 4.** STRING network of genes not expressed in the *sensory system development* gene set (36). This STRING network shows the orthogroups of genes labeled by human gene symbols that are in the gene set that did not contain an *H. symbiolongicarpus* transcript. The key contains the color-coding information for each gene in the STRING networks describing selected biological processes (Gene Ontology) or Annotated Keywords (UniProt).

**Supplementary Figure 5.** Differential gene expression over development in the *transcription factor* gene set. (**A**) Differentially expressed genes over four days of development examining the *transcription factor* gene set derived from the GSEA (33). Each row is a single transcript labeled by the human gene symbols that are present in the same orthogroup as the planula transcript. High expression is yellow and low is light blue. Grey panels contain abbreviated descriptions of gene functions from UniProt (116). (**B-F**) STRING networks showing significantly differentially expressed genes across four days of development where (**B**) contains all DE genes from all four days, and (**C-F**) contains subnetworks of DE genes by day. STRING is a database of known and predicted protein-protein interactions (36). In each interaction map, line thickness indicates the strength of data support. Gene symbols are shaded by biological processes (Gene Ontology) or Annotated Keywords (UniProt). (**G**) Node and edge information for each network that includes the PPI enrichment p-values. PPI enrichment p-values are highest in days 1, when the developmental regulatory network of planula is constructed, and day 4, when the developmental regulatory network underpinning competency of settlement and metamorphosis is established. Expression of transcription factors (TFs) at day 1 expectedly represents a range of functional categories but is largely devoid of the DNA-binding category, which becomes more prominent in later developmental days. We interpret this expression pattern as an initial phase where general TFs, working on a genomic transcriptional landscape set by maternal factors, establish the developmental framework for embryogenesis, followed by later, more specialized TFs with DNA binding capacity. The signature of DNA binding TFs that are differentially expressed in day three and four planulae set the transcriptional stage for metamorphosis.

**Supplementary Figure 6.** Gene trees of orthogroups from Orthofinder of genes used for RNA FISH probes. Red sequences are the target sequences used to create RNA FISH probes and green sequences are the human sequences annotating the orthogroup.

### 1.2 Supplementary Videos

**Supplementary Video 1.** Immunohistochemistry of the nervous system and FMRFamide expression of a whole-mount *H. symbiolongicarpus* planula. Immunohistochemistry staining of a Day 3 larva

(72hpf) where red staining corresponds to acetylated alpha-tubulin of neural cells, magenta corresponds to FMRFamide, a neurotransmitter involved in relaying photosensory information, green corresponds to F-actin in contractile muscle, and blue corresponds to DAPI staining of nuclei.

**Supplementary Video 2.** Immunohistochemistry of the nervous system and FMRFamide expression of an *H. symbiolongicarpus* planula oral end. Immunohistochemistry staining of a Day 3 larva (72hpf) where red staining corresponds to acetylated alpha-tubulin of neural cells, magenta corresponds to RFamide, a neurotransmitter involved in relaying photosensory information, green corresponds to F-actin in contractile muscle, and blue corresponds to DAPI staining of nuclei.
