## Supplementary Figure 2 for "Phototactic preference and its genetic basis in the planulae of the colonial Hydrozoan Hydractinia symbiolongicarpus": Supp_Fig_2.pdf

### Network of Genes that are Not Expressed in Sensory Perception of Light Stimulus Gene set

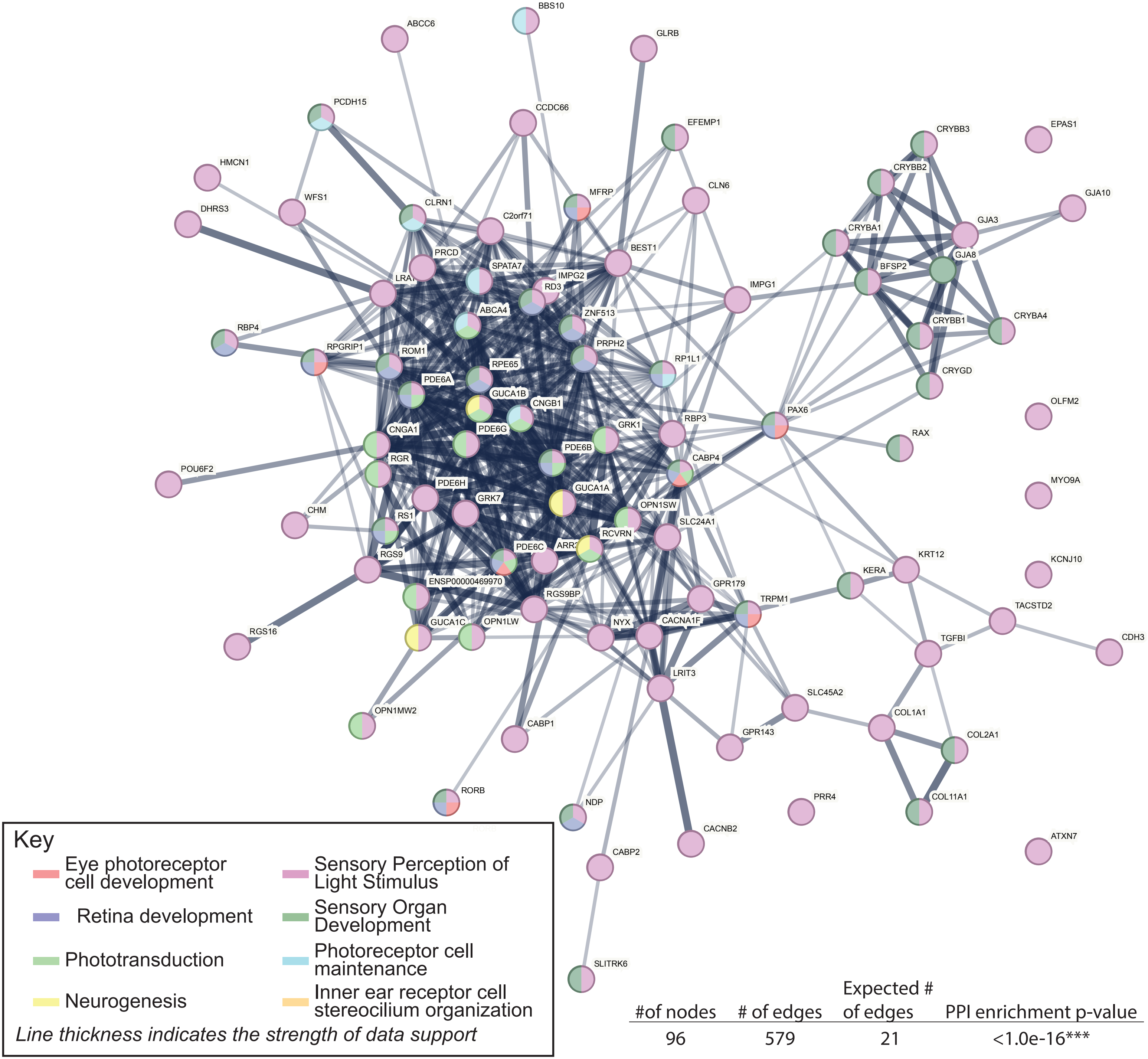


| #of nodes | # of edges | Expected # of edges | PPI enrichment p-value |
|---|---|---|---|
| 96 | 579 | 21 | <1.0e-16*** |
