## Supplementary Figure 4 for "Phototactic preference and its genetic basis in the planulae of the colonial Hydrozoan Hydractinia symbiolongicarpus": Supp_Fig_4.pdf

### Network of Genes that are Not Expressed in Sensory System Development Gene set

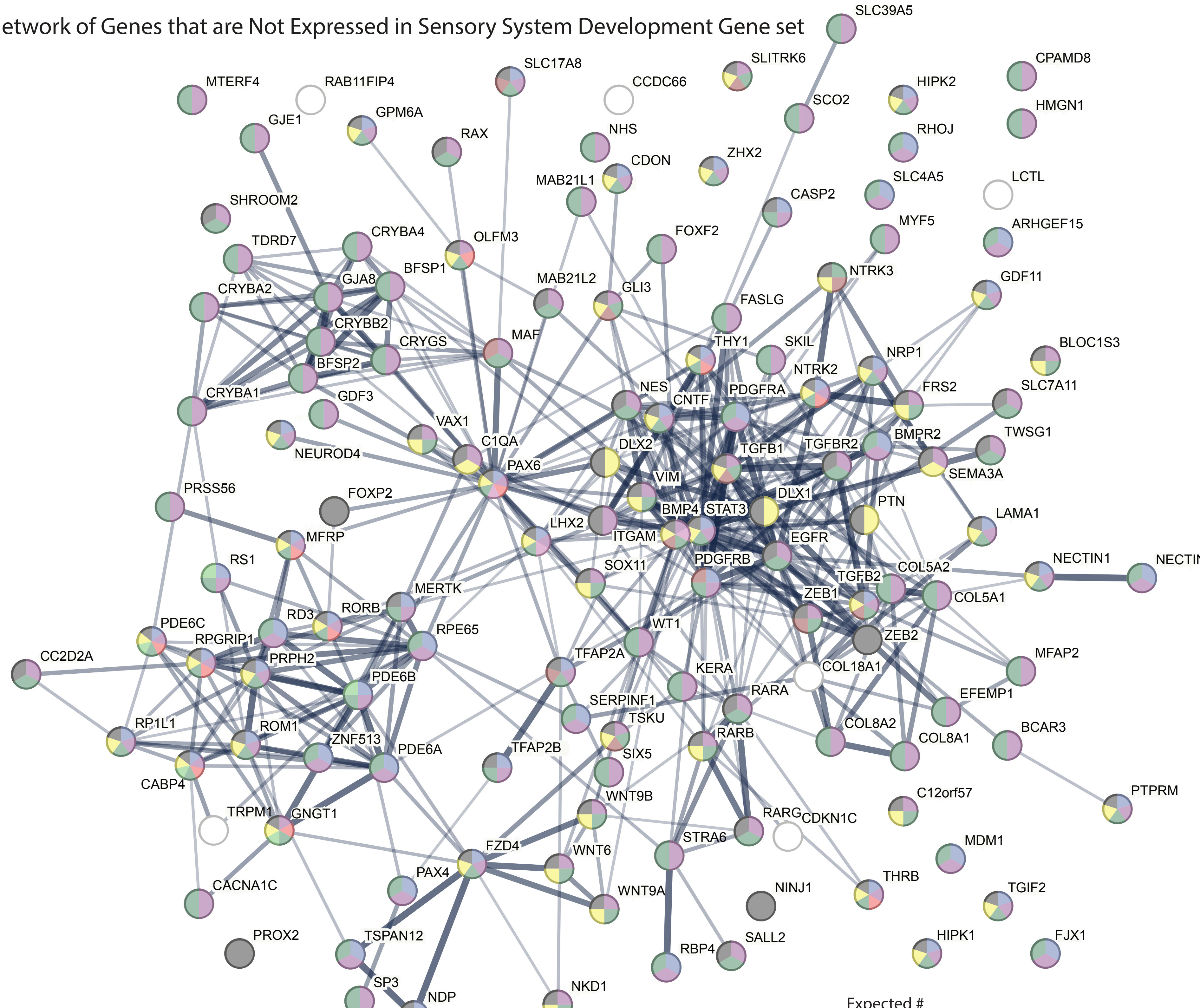


| #of nodes | # of edges | Expected # of edges | PPI enrichment p-value |
|---|---|---|---|
| 140 | 438 | 105 | <1.0e-16*** |

#### Key

- Eye photoreceptor cell development
- Neurogenesis
- Photoreceptor cell maintenance
- Inner Ear Development
- Nervous System Development
- Retina development
- Sensory Perception of Light Stimulus
- Inner ear receptor cell stereocilium organization
- Sensory Perception of Taste
- Retinal Metabolic Process
- Phototransduction
- Sensory Organ Development
- Sensory System Development

*Line thickness indicates the strength of data support*
